# A role for condensin-mediator interaction in mitotic chromosomal organization

**DOI:** 10.1101/2024.06.26.600663

**Authors:** Osamu Iwasaki, Sanki Tashiro, Claire Y. Chung, Tomomi Hayashi, Hideki Tanizawa, Xuebing Wang, Shinya Ohta, Yuko Fujioka, Joseph Han, Gabrielle Tabor, Mikihiro Kawagoe, Ronen Marmorstein, Nobuo N. Noda, Ken-ichi Noma

**Affiliations:** Institute of Molecular Biology, University of Oregon, Eugene, OR 97403, USA; Institute for Genetic Medicine, Hokkaido University, Sapporo 060-0815, Japan; Department of Biochemistry and Biophysics, Perelman School of Medicine, University of Pennsylvania, Philadelphia, Pennsylvania 19104, USA

## Abstract

Eukaryotic genomes are organized by condensin into 3D chromosomal architectures suitable for chromosomal segregation during mitosis. However, molecular mechanisms underlying the condensin-mediated chromosomal organization remain largely unclear. Here, we investigate the role of newly identified interaction between the Cnd1 condensin and Pmc4 mediator subunits in fission yeast, *Schizosaccharomyces pombe*. We develop a condensin mutation, *cnd1-K658E*, that impairs the condensin-mediator interaction and find that this mutation diminishes condensin-mediated chromatin domains during mitosis and causes chromosomal segregation defects. The condensin-mediator interaction is involved in recruiting condensin to highly transcribed genes and mitotically activated genes, the latter of which demarcate condensin-mediated domains. Furthermore, this study predicts that mediator-driven transcription of mitotically activated genes contributes to forming domain boundaries via phase separation. This study provides a novel insight into how genome-wide gene expression during mitosis is transformed into the functional chromosomal architecture suitable for chromosomal segregation.

Graphical Abstract

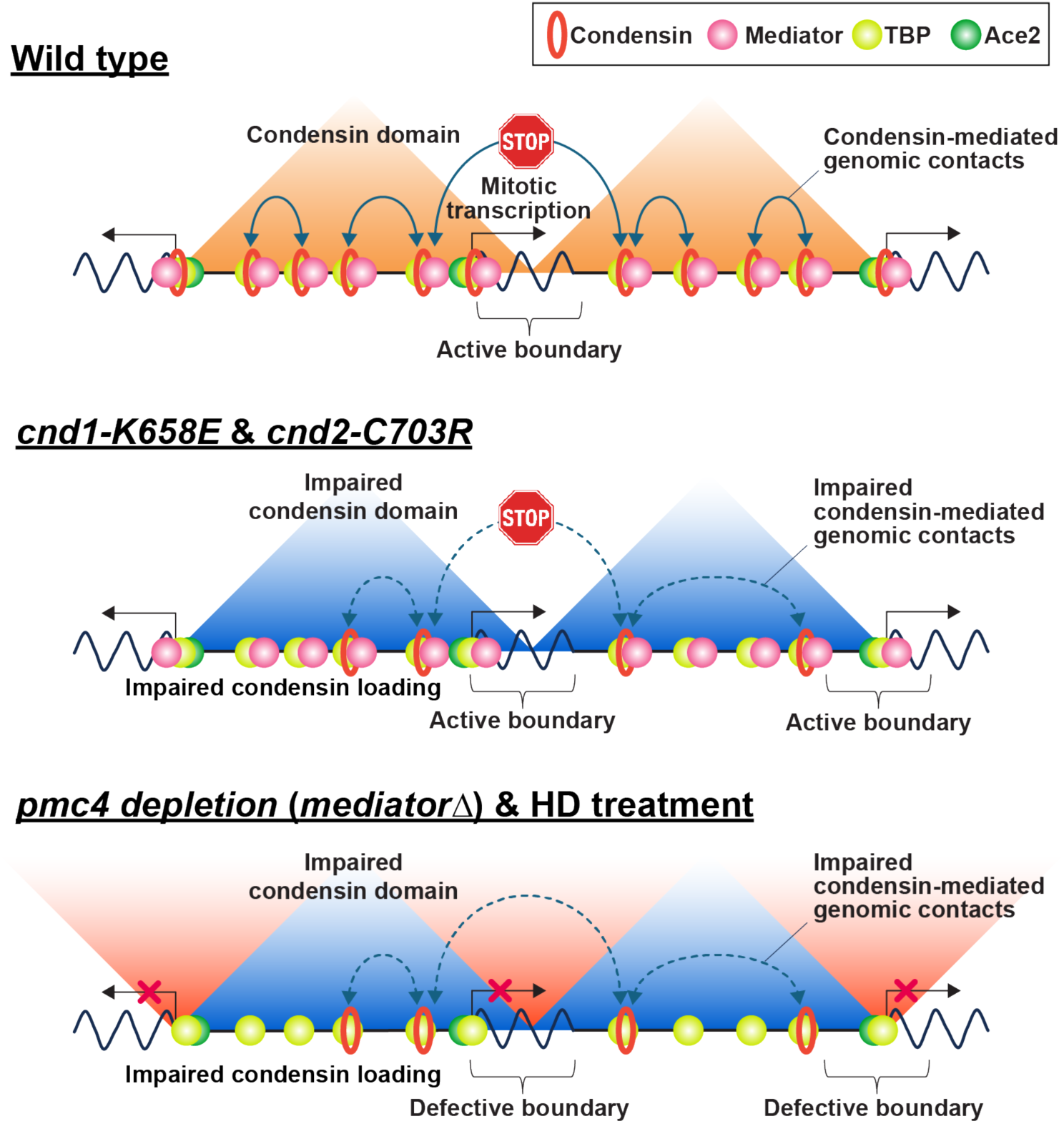

**In Brief:** Mitotic chromosomes are organized by condensin into an architecture suitable for their segregation, although molecular mechanisms underlying how condensin assembles the functional chromosomal architecture remain unclear. This study with the fission yeast model organism shows that the protein interaction between condensin and mediator is involved in chromosomal segregation by forming a mitotic chromosomal architecture. A similar mechanism involving the condensin-mediator interaction is likely conserved in human cells, implying the evolutionary conservation of this mechanism.

**Highlights:**

- The Cnd1 condensin subunit interacts with the Pmc4 mediator component in fission yeast.
- The condensin-mediator interaction is required to recruit condensin to highly transcribed and mitotically activated genes.
- The condensin-mediator interaction is required to form chromatin domains during mitosis.
- Mediator participates in the boundary formation of mitotic chromatin domains.

## INTRODUCTION

The cohesin and condensin complexes consist of Structural Maintenance of Chromosomes (SMC) factors, referred to as SMC complexes (Hirano, 2000; Koshland and Strunnikov, 1996; Nasmyth, 2001; Yanagida, 2009). Cohesin and condensin are evolutionarily conserved among eukaryotes (Hagstrom and Meyer, 2003; Losada and Hirano, 2005) and known to function in sister-chromatid cohesion and mitotic chromosomal condensation, respectively (Freeman et al., 2000; Guacci et al., 1997; Lavoie et al., 2000; Michaelis et al., 1997; Ouspenski et al., 2000; Saka et al., 1994; Tomonaga et al., 2000). Recently, they have also been shown to participate in three-dimensional genome organization (Davidson and Peters, 2021; Hirano, 2012; Noma, 2017; Ong and Corces, 2014; Uhlmann, 2016).

Cohesin forms chromatin contacts and topologically associating domains (TADs), also referred to as cohesin loops and domains (Rao et al., 2017; Rowley and Corces, 2018; Szabo et al., 2019). Cohesin loops are formed via loop extrusion and phase separation mechanisms (Davidson et al., 2019; Fudenberg et al., 2016; Kim et al., 2019; Ryu et al., 2021; Sanborn et al., 2015). TADs are associated with replication timing, and genes within the same TADs along the X chromosome tend to be co-regulated during early differentiation of mouse embryonic stem cells, although understanding the role of TADs in gene regulation requires further investigation (Dixon et al., 2016; Nora et al., 2012; Pope et al., 2014). In this regard, it has been shown that cohesin loss, which eliminates TADs and cohesin loops, has a modest effect on gene regulation across the human genome and that transcriptional elongation disrupts cohesin loops, implying that TADs and loops can simply be reflective of transcriptional processes (Heinz et al., 2018; Rao et al., 2017). That being said, there are some observations that TADs restrict enhancer activity within the domains and prevent developmental genes and oncogenes from ectopic expression, suggesting that TADs become functional in gene regulation at specific developmental stages (Hnisz et al., 2016; Lupianez et al., 2015). Mechanistically, it has been proposed that CTCF forms TAD boundaries by blocking cohesin-mediated loop extrusion (Davidson and Peters, 2021; Rowley and Corces, 2018; Szabo et al., 2019). In fission yeast, it has been shown that TAD boundaries are formed at convergent genes (Kim et al., 2016b; Mizuguchi et al., 2014).

In addition to cohesin, condensin forms chromatin contacts and domains, also referred to as condensin domains in this study (Crane et al., 2015; Gibcus et al., 2018; Kakui et al., 2017; Kim et al., 2016b; Tanizawa et al., 2017; Van Bortle et al., 2014). Condensin-mediated genomic contacts are formed by loop extrusion and diffusion capture mechanisms (Ganji et al., 2018; Kim et al., 2020; Tang et al., 2023). Condensin function is associated with gene regulation (Crane et al., 2015; Dowen et al., 2013; Hocquet et al., 2018; Iwasaki et al., 2019; Lancaster et al., 2021; Li et al., 2015; Swygert et al., 2019). Although it has been shown that CTCF plays a significant role in the formation of TADs and cohesin loops, the factors that participate in the formation of condensin-mediated genomic contacts and domains remain largely unknown.

Our previous study has shown that fission yeast cohesin and condensin form genomic contacts primarily within and beyond 100 kb, respectively (Kim et al., 2016b). Moreover, cohesin and condensin form 30–40 kb and 300 kb–1 Mb chromatin domains, respectively, which are reversely regulated during the cell cycle (Tanizawa et al., 2017). Mechanistically, the Cnd2 condensin subunit interacts with the TBP TATA box-binding protein, Tbp1 in fission yeast, and the disruption of the interaction between Cnd2 and Tbp1 diminishes condensin localization across the genome (Iwasaki et al., 2015). Therefore, it is likely that the interaction between condensin and the TBP general transcription factor participates in condensin loading to gene loci (Noma, 2017). However, how condensin domains are formed, especially regarding the factors involved in forming domain boundaries, remains unclear.

This study demonstrates that the Pmc4 mediator subunit interacts with the Cnd1 condensin subunit and that this interaction is required for the faithful segregation of mitotic chromosomes. We developed a condensin mutation that impairs the condensin-mediator interaction and found that the interaction is required for condensin domain formation. Mechanistically, the condensin-mediator interaction is involved in recruiting condensin to highly transcribed genes and mitotically activated genes present at the boundaries of condensin domains. Mediator is known as a transcriptional regulator/initiator complex (Kornberg, 2005; Malik and Roeder, 2010). This study demonstrates that the functional chromosomal architecture during mitosis is formed by the interaction between the transcriptional machinery, mediator, and the chromosomal compaction factor, condensin. Our observations predict that the mediator-driven transcription of genes present at domain boundaries is responsible for boundary formation via phase separation.

## RESULTS

### Protein interaction between the Cnd1 condensin and Pmc4 mediator subunits

To elucidate how condensin cooperates with other factors to organize the mitotic chromosomal architecture, we sought factors that interact with condensin in fission yeast. For this purpose, we employed a yeast two-hybrid (Y2H) screening approach to identify protein interactions between condensin subunits and factors derived from the cDNA library (the Yeast Genetic Resource Center at Osaka City University). The Y2H screening identified the Pmc4 mediator subunit that potentially interacts with the Cnd1 condensin subunit (**Figure 1A**). It has previously been shown that mediator interacts with cohesin in murine ES cells (Kagey et al., 2010). Therefore, we predicted that mediator would also interact with condensin. To test this, we first examined the Cnd1-Pmc4 interaction in the endogenous fission yeast system by co-immunoprecipitation (co-IP) analysis and detected an interaction (**Figure 1B**). Moreover, fission yeast Cnd1 and Pmc4 proteins were expressed in sf9 insect cells and purified by GST pull-down. Analysis of the fractions from size exclusion chromatography using a Superose 6 column detected a stable and stoichiometric complex between Cnd1 and Pmc4 (**Figure 1C**). These multiple assays consistently pointed to a stable Cnd1-Pmc4 interaction.

**Figure 1.**
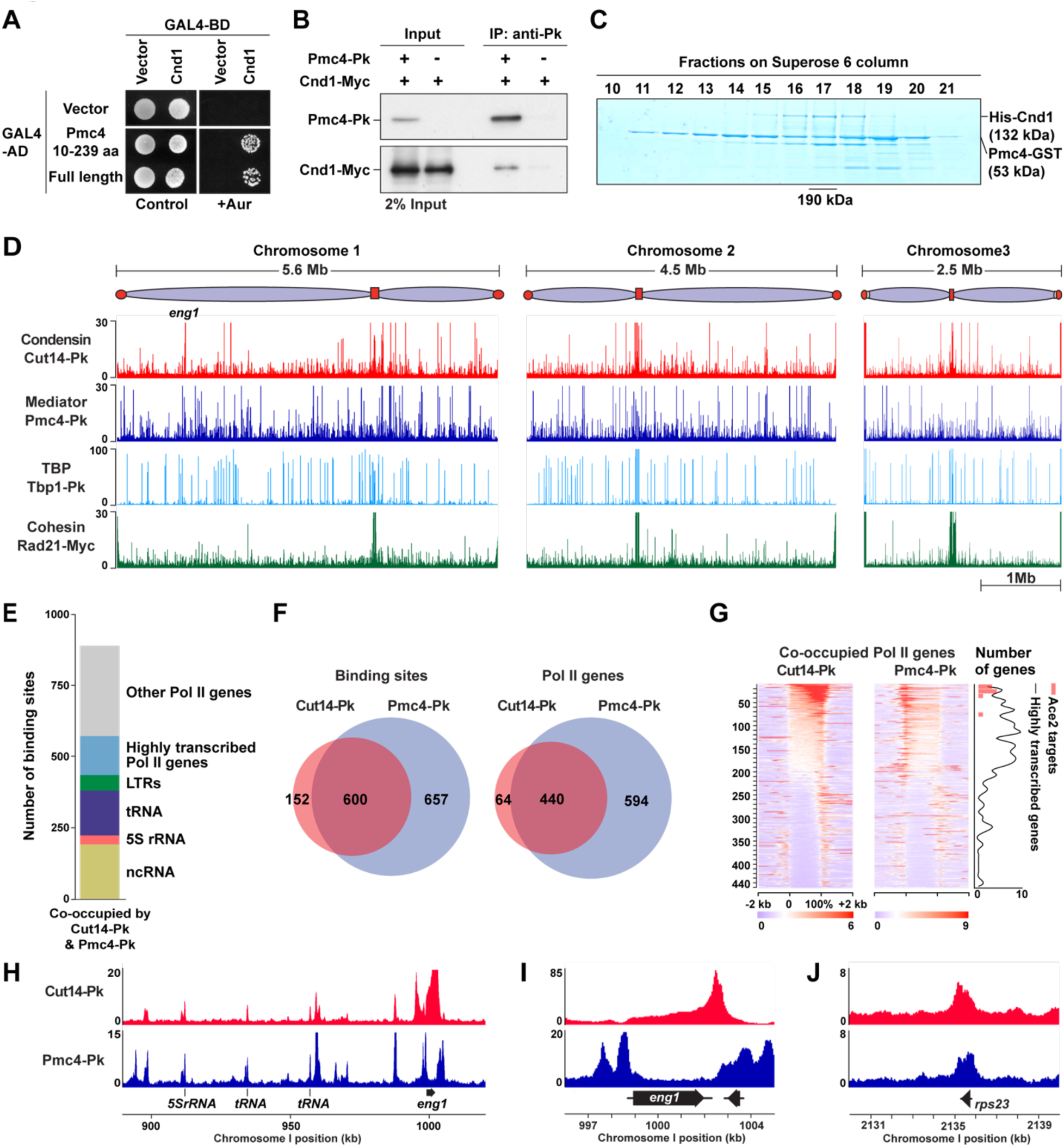
Interaction between condensin and mediator and their co-localization across the fission yeast genome. **(A)** Yeast two-hybrid (Y2H) interaction between Cnd1 condensin and Pmc4 mediator subunits. GAL4-BD-Cnd1 and GAL4-AD-Pmc4 were co-expressed in budding yeast cells. The *pmc4* cDNA fragment encoding 10-239 amino acid (aa) residues of the full-length Pmc4 (239 aa) was initially isolated by Y2H screening. The control plate is SD-Trp-Leu to maintain the plasmids expressing GAL4-BD-Cnd1 and GAL4-AD-Pmc4, and the +Aur plate is SD-Trp-Lue-His-Ade containing 125 ng/ml aureobasidin A to monitor the Y2H interaction. **(B)** Co-IP result showing the interaction between Cnd1-Myc condensin and Pmc4-Pk mediator subunits in fission yeast cells. Cnd1-Myc and Pmc4-Pk were expressed from their endogenous loci. **(C)** GST pull-down analysis on GST-Pmc4 with His-Cnd1. GST-Pmc4 and His-Cnd1 were co-expressed in sf9 insect cells and subjected to GST pull-down, Superose 6 size exclusion chromatography, SDS-PAGE, and Coomassie Brilliant Blue staining. **(D)** Genome-wide distributions of Cut14-Pk (condensin), Pmc4-Pk (mediator), Tbp1-Pk (TBP), and Rad21-Myc (cohesin). **(E)** Summary of binding sites co-occupied by Cut14-Pk and Pmc4-Pk. The top 10% highest transcribed protein-coding genes are categorized as highly transcribed Pol II genes, and the remaining genes are annotated as other Pol II genes. ncRNA stands for non-coding RNA genes. **(F)** Venn diagrams showing the overlap between Cut14-Pk and Pmc4-Pk binding sites (left) and genes (right). ChIP-seq peaks overlapped with ±100 bp regions from transcribed regions were assigned to respective Pol II genes. **(G)** Heatmaps showing Cut14-Pk (left) and Pmc4-Pk (right) distributions at their common target genes (n=440). Pol II genes commonly bound by Cut14-Pk and Pmc4-Pk were ordered from top to bottom based on Cut14-Pk ChIP-seq enrichment. Gene sizes from transcriptional initiation sites (0%) to termination sites (100%) were adjusted to the same length for data representation. Distributions of Ace2 target genes and highly transcribed genes are shown in the right panel. **(H)** Co-localization of Cut14-Pk and Pmc4-Pk at the 130 kb genomic region containing Pol III genes, such as tRNA and 5S rRNA, and the mitotically activated *eng1* gene. **(I-J)** Binding patterns of Cut14-Pk and Pmc4-Pk at the *eng1* **(I)** and *rps23* **(J)** gene loci. The *rps23* is a highly transcribed ribosomal protein gene.

### Co-localization of condensin and mediator at gene regions

If condensin and mediator interact, they may co-localize across the fission yeast genome. To test this possibility, we performed ChIP-seq analyses to map Cut14 condensin and Pmc4 mediator subunits (**Figure 1D**). We found that Cut14 and Pmc4 co-localized at RNA polymerase II-transcribed (Pol II) genes, RNA polymerase III-transcribed (Pol III) genes, such as tRNA and 5S rRNA genes, LTRs, and ncRNA genes (**Figure 1E**). Cut14 and Pmc4 were heavily overlapped at their binding sites (*P* < 1.54 × 10^-760^, hypergeometric distribution test) and Pol II genes (*P* < 1.90 × 10^-690^, hypergeometric distribution test; **Figure 1F**). Among the 440 Pol II genes co-occupied by Cut14 and Pmc4, Cut14 and Pmc4 were preferentially enriched at the 3’ and 5’ ends of genes, respectively (**Figure 1G**). Moreover, Cut14 and Pmc4 were clearly co-localized at Pol II and Pol III genes (**Figure 1H**). Interestingly, high enrichment of Cut14 and Pmc4 was observed at the *eng1* and other mitotically activated genes (**Figure 1G**, **1H**, and **1I**). The Ace2 transcription factor induces transcription of these mitotically activated genes (Petit et al., 2005; Rustici et al., 2004). Cut14 and Pmc4 were also enriched at highly transcribed housekeeping genes, including the *rps23* ribosomal protein gene (**Figure 1J**). These results suggest that condensin and mediator tend to co-localize at highly transcribed genes and mitotically activated Ace2 target genes.

Additionally, we analyzed genome-wide distributions of Tbp1 TATA-box binding protein (TBP) and the Rad21 cohesin subunit because we previously found that Tbp1 interacts with the Cnd2 condensin subunit and that cohesin and condensin both tend to localize at the 3’ end of Pol II genes (Iwasaki et al., 2015; Kim et al., 2016b). We observed that Cut14, Pmc4, Tbp1, and Rad21 were enriched at Pol II genes, Pol III genes, LTRs, and ncRNA genes (**Figure S1A**). Cut14, Pmc4, and Tbp1 were significantly co-localized at their binding sites (*P* < 7.48 × 10^-304^, hypergeometric distribution test) and Pol II genes (*P* < 2.76 × 10^-230^, hypergeometric distribution test), supporting the condensin interactions with TBP and mediator (**Figure S1B**). Consistent with the previous studies, condensin and cohesin were localized at the kinetochore-forming and heterochromatic centromeric regions, respectively (**Figure S1C**) (Iwasaki et al., 2015; Nakazawa et al., 2008; Tomonaga et al., 2000). On the other hand, TBP and mediator were enriched at centromeric tRNA genes, implying that the co-localization between condensin and the transcription-related factors, TBP and mediator, is restricted to gene regions (**Figure S1C**). To further support this finding, Cut14, Pmc4, and Tbp1 were enriched at Ace2 target genes, the top 10% highest transcribed Pol II genes, and Pol III genes (**Figure S1D-F**).

Moreover, we observed a clear difference in condensin and cohesin localizations. Although Rad21 was also localized at the top 10% highest transcribed Pol II genes and Pol III genes, where Cut14, Pmc4, and Tbp1 were also enriched (**Figure S1D-F**), the heatmap illustrating their distributions at all Pol II genes indicated that Cut14, Pmc4, and Tbp1 appeared to tightly co-localize at Pol II genes (n=286), whereas Rad21 was ubiquitously detectable at the 3’ end of many more Pol II genes (**Figure S1G** and **S1H**). Cut14, Pmc4, and Tbp1 were frequently co-localized at highly transcribed Pol II genes and mitotically activated Ace2 target genes (**Figure S1H**). These ChIP-seq and biochemical results collectively suggest that condensin co-localizes with mediator and Tbp1 at gene regions via its interactions.

### Impairment of Cnd1-Pmc4 interaction by *cnd1-K658E* mutation

To investigate the function of the Cnd1-Pmc4 interaction, we set out to generate *cnd1* gene mutations that specifically inhibit the interaction without affecting the condensin complex. We employed a Y2H screening approach combined with PCR-based random mutagenesis and identified the *cnd1-m1* mutations (**Figures 2A**, **S2A**, and **S2B**). The *cnd1-m1* mutations consisted of five point mutations: T584A, K658E, R905H, M1057K, and a non-sense mutation at the Y1136 residue. We dissected these point mutations and found that the *cnd1-K658E* mutation alone could disrupt the Cnd1-Pmc4 interaction (**Figure 2A**). It is important to note that the Cnd1-Cnd2 interaction is known to be the only interaction required for Cnd1 to form the condensin complex (Onn et al., 2007). The Y2H data revealed that the Cnd1-Cnd2 interaction was unaffected by the *cnd1-K658E* mutation, implying that the *cnd1-K658E* mutation impairs the Cnd1-Pmc4 interaction without affecting the condensin complex (**Figure 2A**). This finding was further supported by co-IP analyses using the endogenous fission yeast system, again indicating that the *cnd1-K658E* mutation affects the Cnd1-Pmc4 interaction but not the Cnd1-Cnd2 interaction (**Figure 2B** and **2C**).

**Figure 2.**
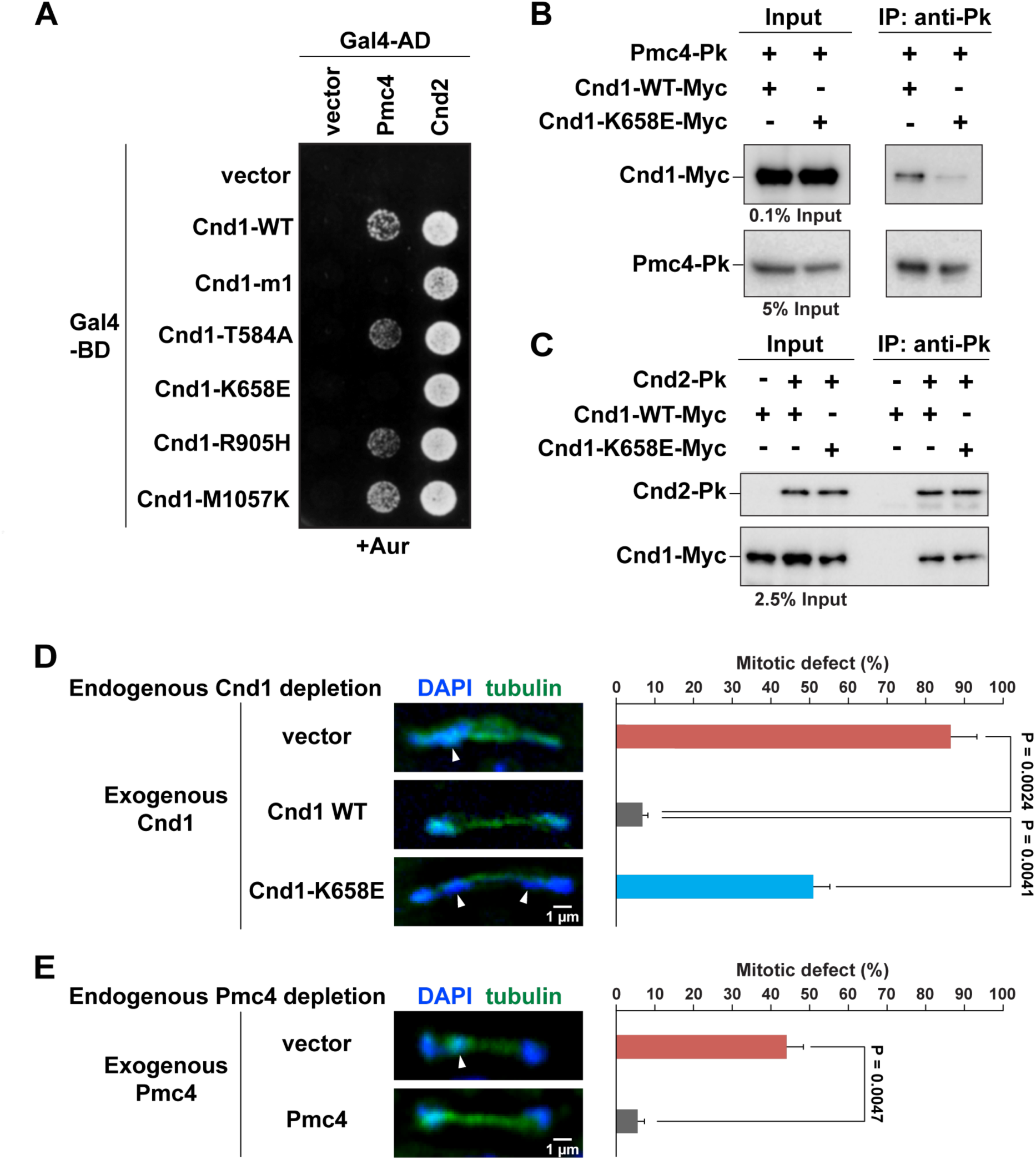
The *cnd1-K658E* condensin mutation impairs the Cnd1-Pmc4 interaction and causes chromosomal segregation defects. **(A)** Y2H analysis to examine the interaction of mutant Cnd1 with the Pmc4 mediator and Cnd2 condensin subunits. Wild-type or mutant Cnd1 fused to GAL4-BD was co-expressed with Pmc4 or Cnd2 fused to GAL4-AD. **(B)** Co-IP analysis investigating the interaction of wild-type and mutant Cnd1-Myc with Pmc4-Pk. **(C)** Co-IP result showing the interactions of wild-type and mutant Cnd1-Myc with Cnd2-Pk. **(D)** Chromosomal segregation defects in cells expressing Cnd1-K658E. Cells were cultured in EMM liquid medium containing thiamine and auxin to induce transcriptional inhibition of the *cnd1* gene and post-translational degradation of endogenous Cnd1 proteins. The wild-type and mutant Cnd1 were expressed from plasmids. The vector represents a negative control without exogenous Cnd1 expression (Cnd1 depletion). IF experiments were performed to visualize spindle microtubules (mitotic marker) and DAPI signals. Mitotic cells with lagging chromosomes (arrowheads) were counted. The experiments were repeated three times. **(E)** Chromosomal segregation defects in cells depleting Pmc4. Cells were cultured in YEA liquid medium containing auxin to deplete the endogenous Pmc4, and the exogenous Pmc4 was expressed from the plasmid. Mitotic defects were examined as described in panel **(D)**. Data in panels **(D)** and **(E)** are represented as mean ± SD.

### Involvement of Cnd1-Pmc4 interaction in chromosomal segregation

We optimized the auxin-inducible degron (AID) system, testing different combinations between degrons (IAA7 and IAA17) and F-box proteins (atTIR1, atAFB2, and osTIR1). For this optimization, the Pmc4 mediator subunit fused to the degron was employed, and we observed that the IAA17 and the two F-box proteins (atAFB2 and osTIR1) worked best (**Figure S2C**). Namely, cells grew well on control plates without auxin, but cell growth was slower on plates with auxin because Pmc4 is essential for cell viability.

To investigate the function of the Cnd1-Pmc4 interaction, we depleted endogenous Cnd1 expression by inhibiting its transcription and targeted protein degradation and expressed wild-type and mutant Cnd1 proteins from plasmids. We observed that cells without the exogenous Cnd1 (vector; Cnd1 depletion condition) did not grow on the EMM-thiamine+Auxin plate, indicating that Cnd1 is indeed depleted (**Figure S2D**). On the other hand, cells grew on the same plate when wild-type and mutant Cnd1 proteins were provided from plasmids, suggesting that the exogenous Cnd1 expression can suppress the depletion of endogenous Cnd1 proteins. Interestingly, the frequency of chromosomal segregation defects in cells expressing Cnd1-K658E was significantly higher than in cells expressing wild-type Cnd1 (**Figure 2D**). We also observed that cells depleting the endogenous Cnd1 and expressing the exogenous Cnd1-K658E grew similarly to those with the wild-type Cnd1, implying that the *cnd1-K658E* mutation is sufficient to cause chromosomal segregation defects to the degree that does not affect the cell viability at the detectable level, offering a valuable genetic tool to investigate function of the condensin-mediator interaction in chromosomal segregation (**Figure S2D**). These results collectively demonstrate that the Cnd1-Pmc4 interaction is required for the fidelity of chromosomal segregation. Moreover, we detected chromosomal segregation defects in cells depleting Pmc4, implying that mediator has a mitotic function in chromosomal segregation through its interaction with condensin (**Figure 2E**).

### Roles of Cnd1-Pmc4 interaction in the formation of condensin-mediated genomic contacts and chromatin domains

We next examined the effects of the *cnd1-K658E* mutation on the 3D chromosomal architecture during mitosis. To this end, we performed in situ Hi-C experiments with mitotic cells depleting endogenous Cnd1 proteins and expressing exogenous the wild-type and mutant Cnd1 (**Figure 3A**). The difference maps and distance curves indicated that Cnd1-K658E expression and Cnd1 depletion decreased genomic contacts approximately from 75 to 800 kb (**Figure 3B** and **3C**). Genomic contacts in this range are mediated by condensin (Tanizawa et al., 2017). The enlarged difference maps consistently revealed that Cnd1-K658E expression and Cnd1 depletion impaired genomic contacts within condensin domains (**Figure 3D**). Contact scores in condensin domains were significantly affected by the condensin mutations (*P* = 1.35 × 10^−23^ and 2.20 × 10^−15^ for Cnd1-K658E expression and Cnd1 depletion, respectively, two-sided paired Student’s *t-*test; **Figure 3E**). These results indicate that the Cnd1-Pmc4 interaction is involved in the formation of condensin-mediated genomic contacts and chromatin domains. In this study, we often employ this 1.9 Mb genomic region as a model locus to examine chromatin domains and boundaries because we previously found that the *eng1* and *SPAC343.20* genes are mitotically activated by Ace2 transcription factor and demarcate condensin domains (Kim et al., 2016b).

**Figure 3.**
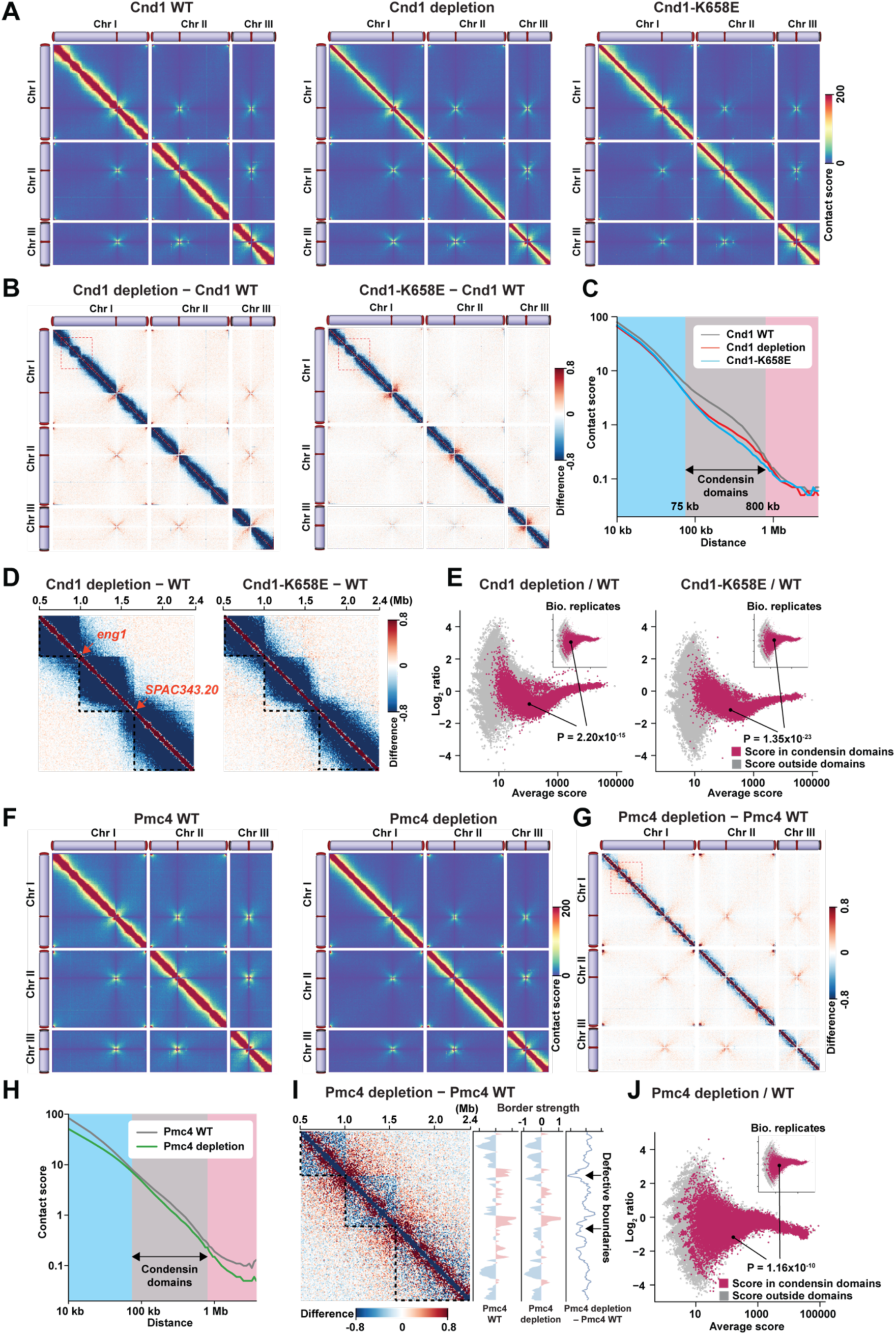
Effects of *cnd1-K658E* condensin mutation and Pmc4 mediator depletion on 3D chromosomal organization during mitosis. (**A**) Genome-wide contact maps from mitotic cells with wild-type Cnd1 expression from the plasmid (Cnd1 WT; left), without exogenous Cnd1 expression (Cnd1 depletion; middle), and with exogenous Cnd1-K658E expression (Cnd1-K658E; right). The endogenous Cnd1 was repressed by culturing cells in EMM medium containing thiamine and auxin. Mitotic cells were prepared by the *cdc25-22* G2/M block-release and harvested at the 60-minute time point after the release. (**B**) Difference maps showing the difference of contact scores between Cnd1 depletion and Cnd1 WT data (left) and between Cnd1-K658E and Cnd1 WT data (right). The Hi-C data in panel **(A)** were used to calculate difference scores. (**C**) Relations between contact scores and genomic distances in the indicated samples. (**D**) Enlarged difference maps for the 1.9 Mb region of chromosome I (Dotted box in panel **B**). The genomic position is shown at the top. The dotted lines indicate previously defined condensin domains (Tanizawa et al., 2017). In this genomic region, the *eng1* and *SPAC343.20* genes function as domain boundaries (Kim et al., 2016b). (**E**) Averages and log_2_ ratios of contact scores between the Cnd1 depletion and Cnd1 WT data (left) and between biological replicates from wild-type mitotic cell cultures (inset). This analysis was performed as previously described (Tanizawa et al., 2017). In brief, genomic contacts within condensin domains (red) were separated from other contacts (gray) and were used in the analysis. A hundred combinations of genomic loci were randomly selected from condensin domains. Log_2_ ratios of contact scores between the Cnd1 depletion and Cnd1 WT data and between the biological replicates were extracted for the selected 100 combinations and subjected to a two-sided paired Student’s *t*-test. Random sampling was repeated 1,000 times, and a median of *P* values was represented. The same analysis was carried out for the Cnd1-K658E and Cnd1 WT data (right). (**F**) Contact maps from mitotic cells with wild-type Pmc4 expression from the plasmid (Pmc4 WT; left) and without exogenous Pmc4 expression (Pmc4 depletion; right). The endogenous Pmc4 was degraded by the AID system. The mitotic cells were prepared using the *cdc25-22* block-release. (**G**) Genome-wide difference map between Pmc4 depletion and Pmc4 WT data. (**H**) Relations between contact scores and genomic distances in Pmc4 WT and Pmc4 depletion data. (**I**) Enlarged difference map between Pmc4 depletion and Pmc4 WT data for the 1.9 Mb region of chromosome I (Dotted box in panel **G**). Border strength in the indicated samples was calculated as described (Tanizawa et al., 2017). Border strength score is, in essence, estimated as upstream and downstream intra-domain contacts divided by inter-domain contacts. The difference in the border strength scores between the samples is shown on the right. Arrows indicate the positions of domain boundaries impaired by Pmc4 depletion. (**J**) Averages and log_2_ ratios of contact scores between the Pmc4 depletion and Pmc4 WT data. The analysis was performed as described in panel (**E)**.

It has previously been shown that Tbp1 interacts with the Cnd2 condensin subunit (Iwasaki et al., 2015). Therefore, we also examined the 3D chromosomal architecture in cells depleting endogenous Cnd2 and expressing the wild-type and mutant Cnd2 (**Figure S3A**). It was shown that the *cnd2-C703R* mutation impairs the Cnd2-Tbp1 interaction without affecting condensin complex formation. We found that Cnd2-C703R expression and Cnd2 depletion significantly diminished condensin domains (**Figure S3A-D**). Of note, the effect of the Cnd2-C703R expression and Cnd2 depletion on domain formation was relatively weak compared with Cnd1-K658E expression and Cnd1 depletion because asynchronous cells were subjected to Hi-C analysis. Nevertheless, these results indicate that the condensin interactions with mediator and TBP are involved in the formation of condensin-mediated genomic contacts and chromatin domains.

### Mediator participates in the formation of condensin-mediated genomic contacts and chromatin domains

To examine the role of mediator in forming the 3D chromosomal architecture during mitosis, we performed in situ Hi-C with mitotic cells degrading Pmc4 proteins using the AID system (**Figure 3F**). The difference map between Pmc4 WT and Pmc4 depletion data and the distance curves revealed that genomic contacts between 75 and 800 kb were impaired by Pmc4 depletion, although the effect was relatively weaker than Cnd1 depletion and Cnd1-K658E expression (**Figure 3G** and **3H**). The enlarged difference map and statistical analysis also indicated that condensin-mediated domains and genomic contacts were significantly disrupted by Pmc4 depletion (**Figure 3I** and **3J**; *P* = 1.16 × 10^−10^, two-sided paired Student’s *t-*test). In addition to the defects in the formation of condensin domains, we observed that Pmc4 depletion enhanced genomic contacts between two adjacent condensin domains and that border strength scores at domain boundaries were reduced by Pmc4 depletion (**Figure 3I**). These results suggest that Pmc4 mediator is involved not only in the formation of condensin-mediated genomic contacts and chromatin domains but also in the assembly of domain boundaries.

Mediator is a large protein complex consisting of approximately 23 subunits in fission yeast (Tsai et al., 2017). Mediator subunits belong to two modules, a core complex and CDK8 regulatory subcomplex, which have distinct implications in different biological pathways (Linder et al., 2008; Malik and Roeder, 2010). We thus expanded our analysis to other mediator subunits and subjected 9 non-essential mediator gene mutants to Hi-C analysis. We found that condensin-mediated genomic contacts and chromatin domains were impaired in *med10Δ*, *med20Δ*, and *pmc3Δ* (*med27Δ*) mutants (**Figure S3E**). However, the effects of these mutations were relatively weaker than Pmc4 depletion, which is potentially because asynchronous cells were subjected to Hi-C analysis for the mediator mutants. We also observed that border strength scores were decreased around the *eng1* gene locus in the same mediator mutants, suggesting that the chromatin boundary at the *eng1* locus was impaired. We also found that gene deletions of CDK8 regulatory subcomplex components, such as *srb10Δ* (*cdk8Δ*) and *srb11Δ*, have little or no effects on condensin domain formation, implicating that the mediator core complex, but not the CDK8 regulatory subcomplex, is involved in the formation of condensin domains during mitosis. These results predict that mediator, but not the Pmc4 subunit alone, participates in forming condensin-mediated genomic contacts and chromatin domains.

### Disruption of condensin domains in the *cnd1-K658E* mutant detected by FISH microscopy

Since the Hi-C analysis reveals that condensin domains were impaired in mitotic cells expressing Cnd1-K658E, we tried to visualize domain disruption by Cnd1-K658E expression using a FISH microscopic approach (**Figure 4A**). For this analysis, we visualized the two condensin domains by FISH and spindle microtubules by IF, allowing us to distinguish interphase and mitotic cells. It is important to note that areas occupied by FISH signals were variable, depending upon fluorescent dye and visualization conditions, and we thus measured the distance between centers of FISH signals as a reproducible parameter to predict the organization of the two condensin domains. We found that, when the wild-type Cnd1 was expressed, centers of the two condensin domains were positioned significantly closer during mitosis than interphase (**Figure 4B**). In contrast, distances between centers of the two condensin domains were significantly extended during mitosis in Cnd1-depleted cells (**Figure 4B**). These results suggest that the shortened distance between centers of the two condensin domains during mitosis observed in cells expressing wild-type Cnd1 depends on condensin. Furthermore, we observed that distances between centers of the two condensin domains also significantly became longer in mitotic cells expressing Cnd1-K658E compared to those expressing wild-type Cnd1 (**Figure 4B**). Distances between centers of the two condensin domains in mitotic cells expressing the Cnd1-K658E were similar to those of Cnd1-depleted mitotic cells, suggesting that the Cnd1-Pmc4 interaction plays a major role in the organization of condensin domains.

**Figure 4.**
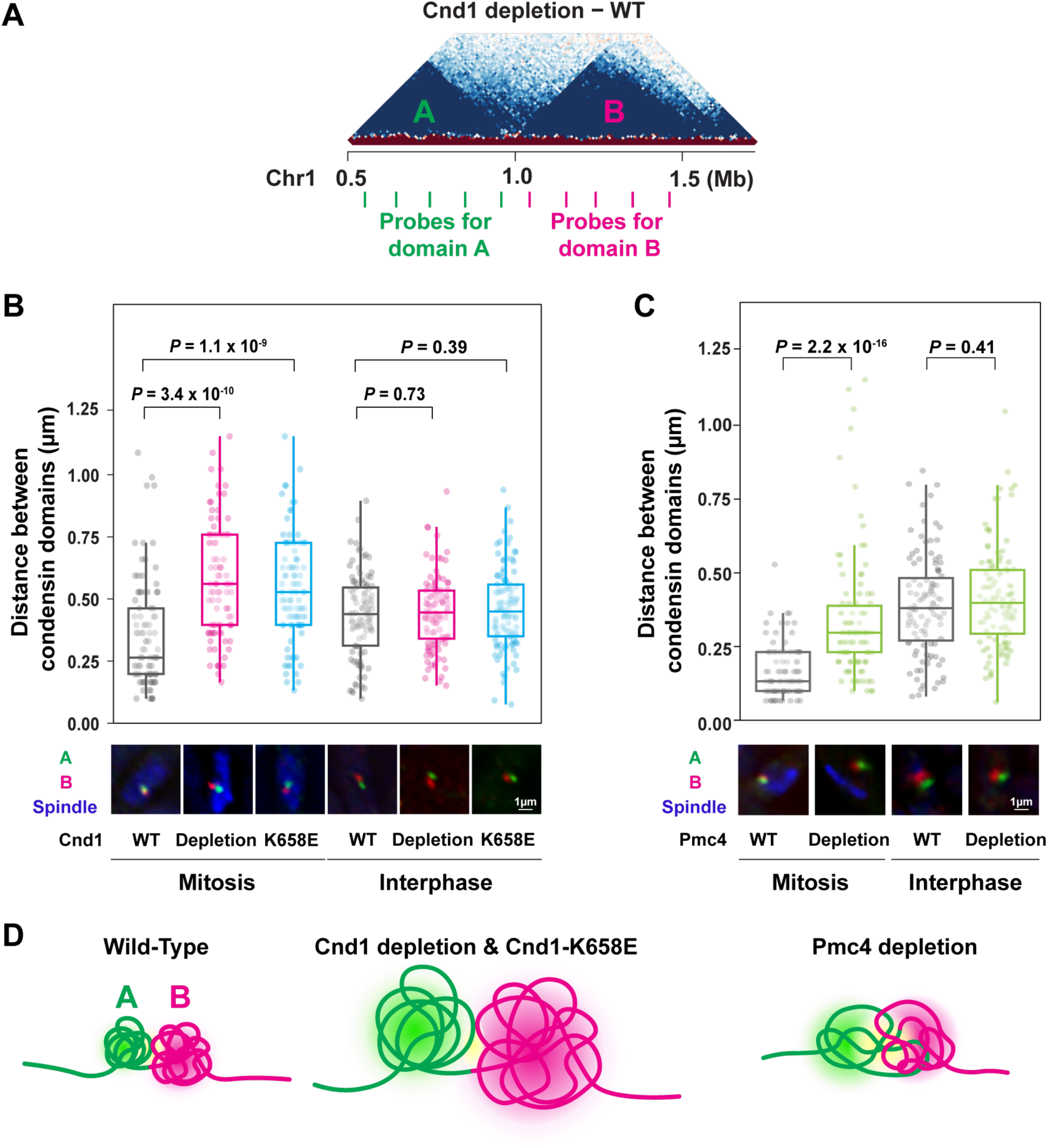
FISH visualization of condensin domains in the *cnd1-K658E* mutant. **(A)** Positions of FISH probes. A and B probes, each consisting of five PCR-derived 5 kb DNA fragments, were used to visualize the indicated condensin domains during mitosis. **(B)** Quantification of distances between centers of two condensin domains in interphase and mitotic cells with wild-type Cnd1 expression from the plasmid (Cnd1 WT), without exogenous Cnd1 expression (Cnd1 depletion), and with exogenous Cnd1-K658E expression (Cnd1-K658E). Typical FISH-IF images were shown underneath, indicating two condensin domains (red and green) and spindle staining (blue; mitotic marker). FISH-stained areas were predicted as condensin domains. **(C)** Quantification of distances between centers of two condensin domains in interphase and mitotic cells with wild-type Pmc4 expression from the plasmid (Pmc4 WT) and without exogenous Pmc4 expression (Pmc4 depletion). The endogenous Pmc4 was removed by the AID system. **(D)** Schematic views of mitotic condensin domains in the indicated conditions. The two adjacent condensin domains are formed during mitosis in wild-type cells, making the two domains closer. In mitotic cells with Cnd1 depletion and Cnd1-K658E mutant, the domains are diminished and thus occupy larger areas, increasing distances between centers of condensin domains. In mitotic cells with Pmc4 depletion, distances between centers of the two domains are shorter than that observed in the Cnd1 depletion and Cnd1-K658E mutant. In panels **(B)** and **(C)**, more than 100 mitotic cells were examined. The central bar represents the median, with boxes indicating the upper and lower quartiles. Whiskers extend to the data points of no more than 1.5× the interquartile range from the box.

Moreover, we designed FISH probes to visualize the two paired genomic loci either within the same domain or spanning the different domains (**Figure S4A**). The visualized two loci are consistently 412 kb away for the intra– and inter-domain FISH probes. We found that when the wild-type Cnd1 was expressed, the intra-domain probes were positioned significantly closer during mitosis than interphase, whereas the inter-domain probes were located closer during mitosis than interphase to a lesser extent compared with the intra-domain probes (**Figure S4B**). The observed difference between the intra– and inter-domain probes represents the domain-based chromosomal compaction during mitosis. Subsequently, we visualized the same paired loci in cells expressing Cnd1-K658E proteins, and the observed difference between the intra– and inter-domain probes was no longer detected, indicating that the domain-based chromosomal compaction is compromised in mitotic cells expressing Cnd1-K658E (**Figure S4B**). These results demonstrate that the Cnd1-Pmc4 interaction promotes domain-based chromosomal compaction during mitosis.

We also visualized the two condensin domains in mitotic cells depleting endogenous Pmc4 proteins with and without exogenous expression of Pmc4. We found that when the wild-type Pmc4 was expressed, centers of the two condensin domains were positioned significantly closer during mitosis than interphase (**Figure 4C**). In addition, distances between the two condensin domains were extended significantly in Pmc4-depleted cells compared to Pmc4-expressing cells (**Figure 4C**). This result suggests that the Pmc4 mediator subunit is involved in the formation of mitotic chromatin domains via its interaction with the Cnd1 condensin subunit (**Figure 4D**).

### The Cnd1-Pmc4 interaction is involved in condensin recruitment

Our results suggested that the Pmc4 mediator and Cnd1 condensin subunits interact and co-localize at highly transcribed genes and mitotically activated genes (**Figure 1**). The disruption of the Cnd1-Pmc4 interaction impaired condensin-mediated genomic contacts and chromatin domains (**Figures 3** and **4**). These results led us to hypothesize that mediator recruits condensin to gene loci. To test this hypothesis, we performed ChIP-seq experiments using mitotic cells depleting the endogenous Cnd1 and expressing wild-type and mutant Cnd1 (**Figure 5A**). We also examined condensin distribution in the cells depleting endogenous Cnd2 and expressing the wild-type and mutant Cnd2 (**Figure 5A**). We observed that the *cnd1-K658E* and *cnd2-C703R* mutations diminished condensin enrichment at highly transcribed Pol II genes, Pol III genes (tRNA and 5S rRNA), and mitotically activated Ace2-target genes (**Figure 5B**-**5F**). These results suggest that mediator and TBP are involved in condensin loading onto gene regions across the genome. Furthermore, we investigated condensin enrichment in mitotic cells depleting Pmc4 and found that Cnd2 condensin enrichment at gene regions was impaired by Pmc4 depletion (**Figure 5A**-**5F**). These results suggest that mediator is located at gene regions to recruit condensin via the Cnd1-Pmc4 interaction.

**Figure 5.**
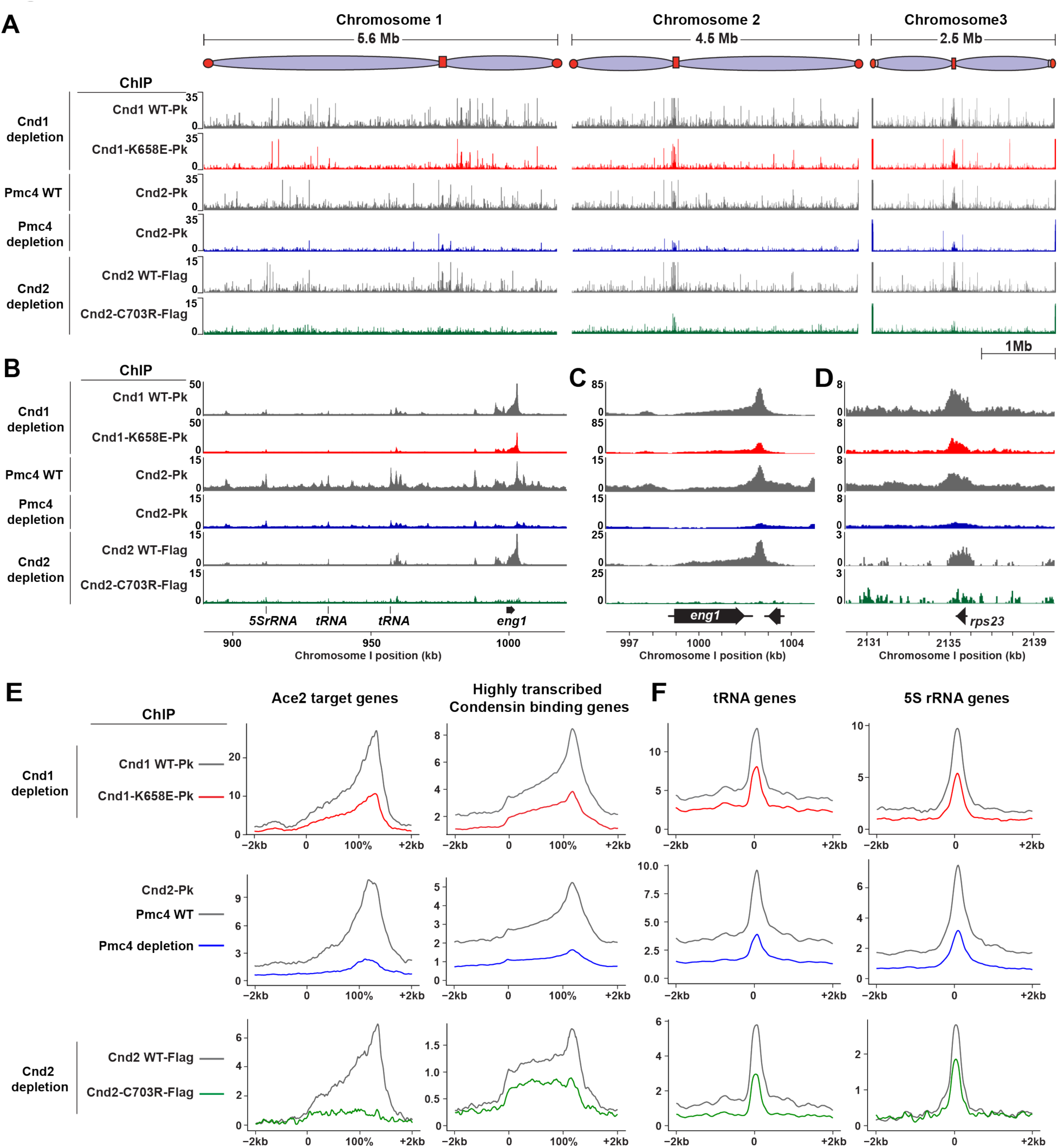
Effects of *cnd1-K658E* mutation on condensin distribution across the genome. **(A)** Genome-wide distributions of condensin were determined by ChIP-seq using mitotic cells with wild-type Cnd1 expression (Cnd1 WT-Pk) and with Cnd1-K658E expression (Cnd1-K658E-Pk) from plasmids. The endogenous Cnd1 was repressed by culturing cells in EMM medium containing thiamine and auxin. Mitotic cells were prepared by the *cdc25-22* G2/M block-release and harvested at a 60-minute time point after the release (top 2 rows). In addition, genome-wide distributions of Cnd2-Pk in cells with wild-type Pmc4 expression from the plasmid (Pmc4 WT) and without exogenous Pmc4 expression (Pmc4 depletion) are shown (middle 2 rows). The endogenous Pmc4 was removed by the AID system. Moreover, genome-wide distributions of the Cnd2-Flag (WT and C703R) expressed from plasmids were determined using cells depleting the endogenous Cnd2 (bottom 2 rows). **(B-D)** Condensin enrichment at the 130 kb genomic region of chromosome I **(B)**, the *eng1* Ace2 target gene **(C)**, and the *rps23* **(D)** highly transcribed gene loci. **(E-F)** Average condensin enrichment at Ace2 target genes and highly transcribed genes **(E)**, and at tRNA and 5S rRNA genes **(F)** in the indicated samples.

### Opposing effects of Pmc4 depletion on highly transcribed genes and mitotically activated genes

To explore the potential role of the condensin-mediator interaction in transcription, we examined global expression profiles in cells expressing Cnd1-K658E. For this purpose, we employed the nascent RNA labeling as described previously (Mahat et al., 2016), followed by deep sequencing (nascent RNA-seq) to determine nascent transcripts. We found that Cnd1-K658E expression and Cnd1 depletion had a modest effect on transcription (**Figure S5A**-**S5E**). We also carried out nascent RNA-seq experiments using Pmc4-depleted cells (**Figure S5A-S5D**) and found that *eng1* transcription was almost completely abolished by Pmc4 depletion (**Figure S5C**). Among 87 significantly down-regulated genes, 8 were Ace2 target genes (**Figure S5F**). We confirmed that condensin genes were not significantly affected by Pmc4 depletion.

Next, we compared the nascent RNA-seq expression profiles with ChIP-seq enrichment of Cut14-Pk condensin, Pmc4-Pk mediator, and Tbp1-Pk. We observed that gene expression levels were tightly correlated with ChIP enrichment of these factors, as nascent RNA-seq strong signals coincided with high ChIP enrichment (**Figure S5G**). Intriguingly, genes with the highest enrichment of these factors were typically Ace2-targeted mitotically activated genes, which were down-regulated by Pmc4 depletion (**Figure S5H** and **S5I**). Moreover, approximately 300 genes with high enrichment of condensin, mediator, and TBP were generally highly transcribed (**Figure S5H**). Those 300 highly transcribed genes were often slightly up-regulated by Pmc4 depletion (**Figure S5H** and **S5J**). Based on these observations, we predict that mediator binding is inhibitory for the excess expression of highly transcribed genes but promotes the expression of mitotically activated Ace2 target genes present at domain boundaries.

### Involvement of phase separation in the formation of mitotic chromosomal architecture

Human Med1 in the mediator complex contains intrinsically disordered regions (IDRs) and is known to form droplets via liquid-liquid phase separation (Cho et al., 2018; Sabari et al., 2018), and the droplet formation by Med1 is connected to transcriptional initiation (Boija et al., 2018). In fission yeast, we noticed that Med1 was not annotated with IDRs. Instead, Pmc4 is predicted to have a disordered region at the C-terminal end (**Figure S6A**). Therefore, we hypothesized that Pmc4 might form liquid droplets. To test this hypothesis, we first purified Pmc4 proteins expressed in *Escherichia coli* and observed that Pmc4 formed droplets in conditions containing more than 5% PEG (**Figure S6B** and **S6C**). Moreover, we examined Pmc4 proteins in fission yeast cells and observed that the treatment with 1,6-hexanediol (HD), which is known to suppress liquid-liquid phase separation, disrupted the focal localization of Pmc4 proteins in nuclei, whereas Rbp1 Pol II foci were retained in nuclei at the same conditions (**Figure 6A**). These results imply that Pmc4 proteins tend to form droplets via phase separation. To gain more insights into the subnuclear localization of mediator and condensin, we applied Zeiss Airyscan super-resolution microscopy to mitotic nuclei and observed that condensin forms fiber-like structures along mitotic chromosomes (**Figure 6B**). On the other hand, mediator was observed as circular spots on mitotic chromosomes (**Figure 6B**). These results collectively suggest that mediator forms transcription-related condensates on mitotic chromosomes and contributes to the formation of condensin-mediated chromatin domains.

**Figure 6.**
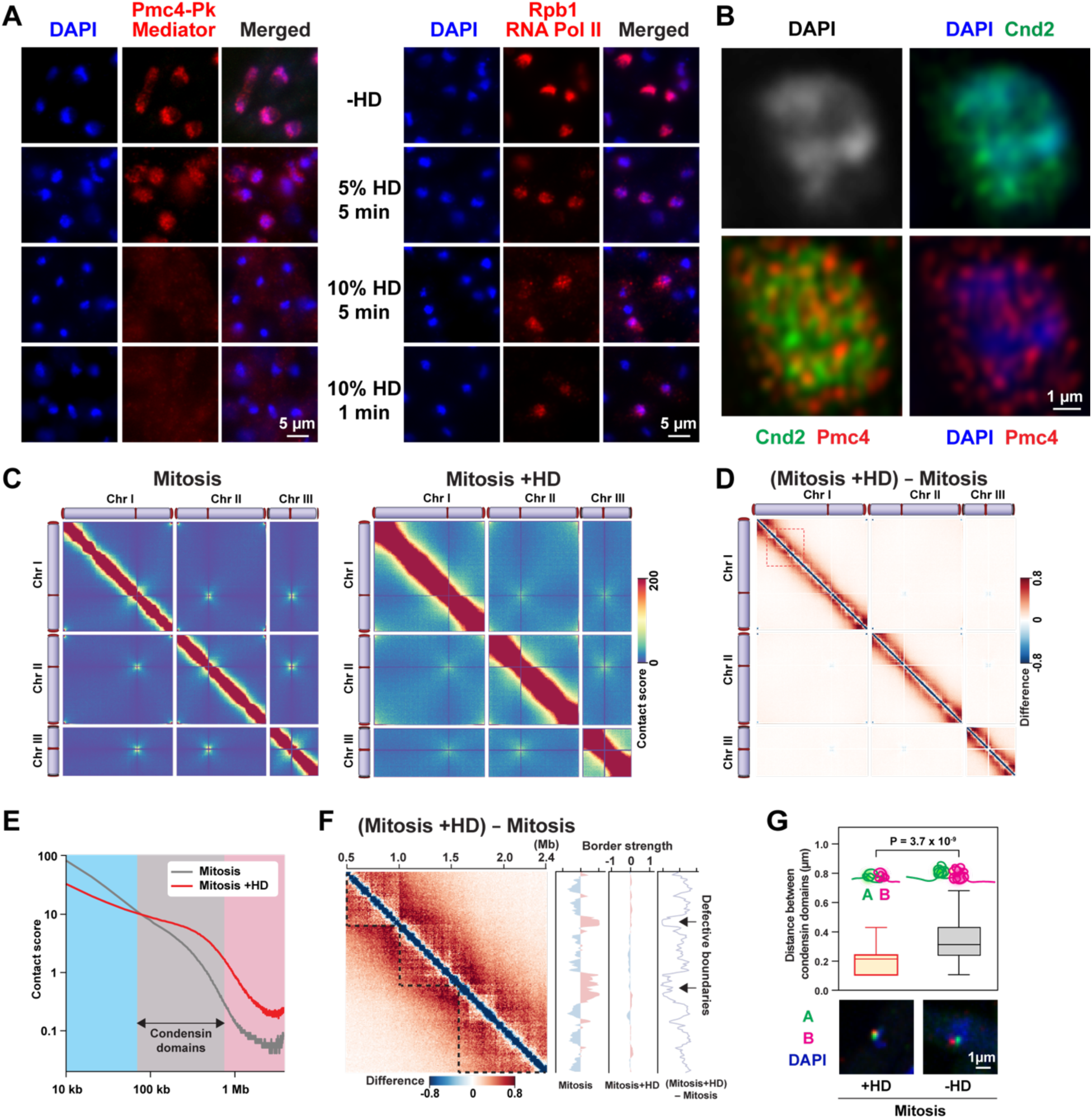
Potential involvement of phase separation in mitotic chromosomal organization. **(A)** Effects of HD treatment on Pmc4 localization. Fission yeast cells were treated with 5 or 10% HD for 1 or 5 minutes and subjected to IF visualization of Pmc4-Pk and Rpb1 (RNA Pol II subunit). **(B)** Subnuclear localization of Pmc4 and Cnd2 visualized using Zeiss Airyscan super-resolution microscope. **(C)** Genome-wide contact maps from mitotic cells treated (right) and untreated (left) with HD. Mitotic cells were prepared by the *cdc25-22* G2/M block-release and treated with 10% HD for 5 minutes. **(D)** Difference maps showing the difference of contact scores between ±HD mitotic samples. **(E)** Relations between contact scores and genomic distances in the ±HD mitotic samples. **(F)** Enlarged difference map between ±HD mitotic samples for the 1.9 Mb region of chromosome I (Dotted box in panel **D**). Border strength scores in the indicated samples and their difference was estimated as described in Figure 3I. **(G)** Quantification of distances between centers of two condensin domains in HD-treated mitotic cells. Mitotic cells were prepared by the *cdc25-22* G2/M block-release. Typical FISH images were shown underneath, indicating the two condensin domains (red and green) and DAPI signals (blue). FISH experiments and data analysis were carried out as described in Figure 4.

Therefore, we examined how the HD treatment affects the mitotic chromosomal architecture. To this end, we again employed an in situ Hi-C approach (**Figure 6C**). The difference map between ±HD mitotic samples and distance curves revealed that long-range genomic contacts (> approximately 75 kb) were elevated in HD-treated cells (**Figure 6D** and **6E**). We noticed that genomic contacts within condensin domains were relatively unaffected by the HD treatment, whereas genomic contacts spanning condensin domains were promoted (**Figure 6F**). This result indicates that the HD treatment compromises domain boundaries, but chromosomal compaction proceeds without forming mitotic chromatin domains. Furthermore, we employed a FISH-IF approach to visualize the two condensin domains in mitotic cells with and without HD treatment. Interestingly, distances between centers of the two condensin domains during mitosis became shorter in HD-treated cells than those without HD treatment, implying that phase separation is involved in separating mitotic chromatin domains (**Figure 6G**). The Hi-C data and microscopic observations suggest that phase separation promotes the formation of mitotic chromatin boundaries (see Discussion).

### Potential conservation of condensin-mediator interaction in human cells

Since condensin and mediator are conserved in fission yeast and humans, the condensin-mediator interaction might also be conserved in human cells. To test this possibility, we employed the Y2H approach to test the interaction. There are two condensin complexes, condensin I and II, in human cells, and we tested the interactions of Med4 (human Pmc4) with CAP-D2 and CAP-D3, human Cnd1 in the condensin I and II complexes, respectively (**Figure S6D**). Interestingly, we observed the Med4 interaction with CAP-D3 (condensin II subunit) but not with CAP-D2 (condensin I subunit). Moreover, the interaction was disrupted by the *CAP-D3-V628E* mutation, which is cognate with the fission yeast *cnd1-K658E* mutation, and by the *CAP-D3-Δ613-635* deletion covering the point mutation (**Figure S6E**). In addition, mitotic defects, such as misaligned chromosomes during metaphase and micronuclei, were detected in human U2OS cells expressing the CAP-D3-V628E, and frequencies of mitotic defects were significantly higher in mitotic cells expressing CAP-D3-V628E than those expressing the wild-type counterpart (**Figure S6F** and **S6G**). Therefore, we speculate that the mitotic mechanism similar to fission yeast chromosomal organization involving the condensin-mediator interaction might function in human cells.

## DISCUSSION

### Condensin loading via its interactions with mediator and TBP

Our ChIP-seq data indicate that condensin tightly co-localizes with the Pmc4 mediator subunit and Tbp1 (fission yeast TBP) at highly transcribed genes and mitotically activated Ace2 target genes (**Figure S1**). Condensin accumulates at the 3’ end of its target genes, whereas peaks of Pmc4 and Tbp1 are positioned at gene promoters. Mechanistically, the *cnd1-K658E* and *cnd2-C703R* condensin mutations inhibit the condensin interactions with Pmc4 and Tbp1, respectively, and diminish condensin localization at its target genes (**Figure 5**) (Iwasaki et al., 2015). Therefore, mediator and TBP likely recruit condensin to promoter regions, and transcription relocates condensin along gene bodies to the 3’ end of genes. In this regard, cohesin relocation by transcription was reported previously (Lengronne et al., 2004). Currently, the cooperative action between the condensin interactions with Pmc4 and Tbp1 remains unclear, but TBP in the TFIID complex and Med4 (Pmc4) in the mediator complex are positioned in proximity to where Pol II transcription initiates (Richter et al., 2022; Schilbach et al., 2017). Therefore, we speculate that the Cnd1-Pmc4 and Cnd2-Tbp1 interactions can facilitate the condensin recruitment to gene promoters.

### Mitotic chromatin boundaries formed by mediator-driven transcription

In the *cnd2-C703R* and *cnd1-K658E* mutants, condensin-mediated genomic contacts within mitotic chromatin domains are compromised (**Figures 3D** and **S3C**; see **Graphical Abstract**). This is probably because condensin localization is diminished at highly transcribed genes and mitotically activated across the genome (**Figure 5**). In contrast, inter-domain genomic contacts are elevated by Pmc4 mediator depletion, while condensin-mediated genomic contacts and mitotic chromatin domains are impaired (**Figure 3I**). Condensin-mediated genomic contacts and chromatin domains are impaired because the Pmc4-dependent condensin loading is compromised, but residual condensin at gene regions can mediate inter-domain genomic contacts in Pmc4-depleted cells (**Figure 5**). If condensin localization is sufficient to form chromatin boundaries, similar Hi-C data should be observed between the *cnd1-CK658E* mutant and Pmc4-depleted cells, but that was not the case (**Figure 3D** and **3I**). Moreover, Pmc4 depletion abolishes the expression of Ace2 target genes present at chromatin boundaries and compromises boundaries (**Figure S5C** and **S5F**). Therefore, we propose that mediator-driven transcription is associated with the boundary formation of mitotic chromatin domains (**Graphical Abstract**).

### Involvement of phase separation in mitotic chromosomal organization

Mediator was initially identified as part of the RNA polymerase II holoenzyme complex and is known to be involved in transcriptional initiation and repression (Ding et al., 2008; Kim et al., 1994). Mediator is also involved in transcriptional regulation in fission yeast (Linder et al., 2008). It has recently been proposed that mediator participates in transcription via phase separation (Cho et al., 2018; Sabari et al., 2018). This stimulated us to investigate the potential involvement of phase separation in the mitotic chromosomal organization involving mediator. In human cells, Med1 is known to form liquid droplets, but Med1 in fission yeast is not annotated as an IDR-containing protein. On the other hand, Pmc4 (fission yeast Med4) contains an IDR at the C-terminal end and is predicted to form droplets. Indeed, our in vitro data suggest that Pmc4 potentially forms liquid droplets. Moreover, HD treatment disrupts mediator localization in the nucleus, suggesting that fission yeast mediator can be regulated by phase separation. Also, we find that the mitotic chromosomal structure is compromised in HD-treated cells, where chromosomal compaction occurs without forming chromatin domains (**Figure 6F**). In addition, as discussed above, Pmc4 mediator depletion compromises domain boundaries. These results collectively suggest that mediator-driven transcription involves phase separation, as observed in other organisms, and promotes boundary formation of mitotic chromatin domains.

Taking all these results together, our mechanistic view is the following: (1) Mediator and TBP recruit condensin to highly transcribed genes and mitotically activated Ace2 target genes; (2) Condensin at gene regions mediates genomic contacts; (3) Mediator activates transcription of Ace2 target genes via phase separation; (4) Transcription of Ace2 target genes demarcates condensin-mediated genomic contacts; and (5) Domain-based chromosomal compaction facilitates proper chromosomal segregation.

## EXPERIMENTAL PROCEDURES

### Fission yeast strains and culture conditions

Using a PCR-based module method, Cnd1, Cnd2, and Pmc4 were tagged with Myc, Flag, and Pk at the C terminus (Bahler et al., 1998; Gadaleta et al., 2013). Genes encoding the tagged proteins were controlled by their own promoters and expressed from their endogenous loci. Other DNA constructs for gene deletions, point mutations, and degron-based targeted protein degradation (nmt81-Cnd1-IAA17 and Pmc4-IAA7) were generated by PCR and introduced to fission yeast cells by the standard chemical transformation. Other strains were constructed by conventional genetic crosses. Fission yeast (*Schizosaccharomyces pombe*) cells were cultured in yeast-extract adenine (YEA) or Edinburgh minimal medium (EMM).

### Auxin-inducible degron (AID) approach for targeted protein degradation

The AID system was employed as previously described (Kanke et al., 2011; Nishimura et al., 2009). The Cnd2-AID system was employed as previously described (Iwasaki et al., 2015). Skp1 fused to the F-box proteins (atAFB2, atTIR1, and osTIR1), nuclear localization signal (NLS), and 9×Myc was under the control of the *adh1* promoter and expressed from the *ade6* locus. The Cnd1 fused to the IAA17 degron was controlled by the *nmt81* promoter and expressed from the endogenous *cnd1* locus. Wild-type and mutant Cnd1 proteins under the control of the endogenous *cnd1* promoter were expressed from LEU2-based plasmids. Cells were cultured in EMM medium containing 15 μM thiamine and 0.5 mM auxin [1-Naphthaleneacetic acid (Sigma)] to induce transcriptional inhibition of the *cnd1* gene and post-translational degradation of endogenous Cnd1 proteins. Pmc4 fused to the IAA7 degron was expressed from the endogenous *pmc4* gene locus. Cells were cultured with YEA medium containing 0.5 mM auxin to induce the Pmc4 degradation.

### Generation of *cnd1* mutations using yeast two-hybrid (Y2H) approach

The schematic procedure is shown in **Figure S2A**. A genetic screen was performed using the Matchmaker Gold Y2H system (Clontech) to generate *cnd1* mutations that disrupt the Cnd1-Pmc4 interaction without affecting the condensin complex. To this end, the *cnd1^+^* gene was first amplified by error-prone PCR (McCullum et al., 2010), and the mutated *cnd1* gene was cloned into pGBKT7 carrying the *TRP1* marker gene, allowing Mat α budding yeast cells to express GAL4-BD fused to the mutant Cnd1 (GAL4-BD-mutant Cnd1). The pGADT7 derivatives that express GAL4-AD fused to either Pmc4 or Cnd2 (GAL4-AD-Pmc4 or GAL4-AD-Cnd2) in addition to the *LEU2* marker gene were independently transformed into the Mat a Y2HGold budding yeast strain (Clontech). After crossing Mat α strain with Mat a strain on YEA plates, diploid cells carrying pGBKT7 and pGADT7 derivatives were selected on Synthetic Defined (SD)-Trp (tryptophan) –Leu (leucine) plates by replica plating. The protein interactions were examined by monitoring cell growth on SD-Trp-Leu-His (histidine)-Ade (adenine) plates containing 125 ng/ml aureobasidin A, reflecting the expression of the *HIS3*, *ADE2*, and *AUR1-C* markers. More than 3,000 diploid colonies that grew on SD-Trp-Leu plates were prepared, and the GAL4-BD-Cnd1-m1 clone was identified. Note that diploid cells carrying pGBKT7 GAL4-BD-Cnd1-m1 and pGADT7 GAL4-AD-Pmc4 did not grow on SD-Trp-Leu-His-Ade plate containing aureobasidin A, but cells carrying pGBKT7 GAL4-BD-Cnd1-m1 and pGADT7 GAL4-AD-Cnd2 could grow on the selection plate. The *cnd1-m1* mutation was determined by Sanger sequencing. The *cnd1-m1* mutation carrying several point mutations was further dissected by generating pGBKT7 GAL4-BD-mutant Cnd1 carrying different point mutations individually (**Figure S2B**).

### Co-immunoprecipitation (co-IP)

Co-IP experiments were performed as described (Iwasaki et al., 2015). Logarithmically growing fission yeast cells in 10 ml YEA (1 x 10^8^ cells at OD_595_ = 0.5) were collected and washed with 1 ml cold H_2_O. Cell pellets were suspended in 500 μl of IP buffer 1 [50 mM HEPES (pH 7.6), 125 mM KCl, 0.1% NP-40, 20% glycerol, 1 mM EDTA, 1 mM PMSF, and Complete protease inhibitor cocktail (Sigma)], and disrupted by Mini-Beadbeater-16 (BioSpec Products). The soluble fraction was collected by centrifugation, and MgCl_2_ (final 5 mM) and 5 units of RQ1 RNase-free DNase I (Promega) were added. After incubating at 37°C for 30 minutes, the DNase I reaction was terminated by adding EDTA (final 10 mM). After the DNase I treatment, the soluble fraction was mixed with Dynabeads Protein G (Life Technologies) coupled with mouse monoclonal anti-Pk (BioRad) antibodies and incubated at 4°C for 2 hours. After washing the beads 5 times for 5 minutes with 800 μl of IP buffer 2 [50 mM HEPES (pH 7.6), 125 mM KCl, 0.2% NP-40, 20% glycerol, 1 mM EDTA, and 1 mM PMSF], proteins were eluted by boiling with 10 μl 1× Laemmli sample buffer.

### GST pull-down

The GST pull-down experiment was performed as previously described (Ricketts et al., 2015). Full-length Pmc4 and Cnd1 were amplified PCR and cloned into pFastBacGST (Fisher Scientific) and pFastBacHTB (Fisher Scientific), respectively. GST-Pmc4 and His-Cnd1 proteins were co-expressed in baculovirus-infected sf9 insect cells and purified by GST pull-down. The GST pull-down sample was subjected to size exclusion chromatography using the Superose 6 column (Sigma), and the fractions were separated by SDS-PAGE, and proteins were stained with Coomassie Brilliant Blue G-250 (Sigma).

### Immunofluorescence (IF) microscopy

Fission yeast cells (1 x 10^8^ cells at OD_595_ = 0.5) were fixed with 3% paraformaldehyde (pFA) at 26°C for 10 minutes, and the fixation was quenched by incubating with glycine (final 0.25 M) for 5 minutes. The fixed cells were washed with PEM buffer [100 mM PIPES (pH 6.9), 1 mM MgCl_2_, and 1 mM EGTA], incubated with PEMS buffer [100 mM PIPES (pH 6.9), 1mM MgCl_2_, 1 mM EGTA, and 1 M sorbitol] at room temperature for 5 minutes, and permeabilized with 1 mg/mL Zymolyase 100 T (Seikagaku) in PEMS buffer at 37°C for 30 minutes. After incubating with PEMS buffer containing 1% Triton X-100 at room temperature for 1 minute and washing twice with PEM buffer, the permeabilized cells were treated with PEMBAL buffer [100 mM PIPES (pH 6.9), 1 mM MgCl_2_, 1 mM EGTA, 1 % BSA, and 0.1 M L-lysine] at room temperature for 1 hour with a gentle rotation. Subsequently, the mouse monoclonal antibodies [anti-Myc (Takara Bio), anti-Pk (BioRad), anti-Pol II antibody CTD4H8 (Abcam), anti-tubulin TAT-1 (Woods et al., 1989)], and the rabbit polyclonal antibodies [anti-Cnd2 (Cosmo Bio USA)] were added at 1/1000 dilution in PEMBAL solution, and the cells were incubated at room temperature overnight. After washing with PEMBAL buffer 3 times, the secondary antibodies [Alexa Flour 488-conjugated anti-rabbit IgG (Life Technologies) and Cy3-conjugated anti-mouse IgG (Jackson ImmunoResearch)] were added at 1/1000 dilution in PEMBAL buffer, and the cells were incubated at room temperature for 3 hours. The cells were washed 3 times with PEMBAL buffer by rotating at room temperature for 15 minutes. Prior to the microscopic investigation, DNA was stained with 1 μg/mL DAPI (Fisher) in PBS solution, and the stained cells were mounted on a coverslip with antifade solution [1 mg/ml p-phenylenediamine, 90% glycerol, and 100 mM Tris-HCl (pH 8.0)]. The fluorescent signals were detected using a DeltaVision deconvolution microscope (GE Healthcare). IF experiments were repeated at least three times, and more than 30 mitotic cells were analyzed for respective samples unless otherwise noted. Super-resolution microscopic images were acquired on a Zeiss LSM 880 with Airyscan running ZEN v. 2.3 (Zeiss). Images were acquired using the 405 nm diode laser for excitation of DAPI, the 488 nm line of an argon laser for excitation of AlexaFluor 488, and the 561 DPSS laser for excitation of Cy3. Three-dimensional Airyscan processing was done using ZEN 2.3 with “Auto” filter strength.

### Fluorescence in situ hybridization (FISH) microscopy

FISH probes were prepared using PCR-amplified DNA fragments as described previously (Iwasaki et al., 2016). To detect a specific condensin domain in the fission yeast chromosome 1, five pairs of primers were designed to amplify 5 kb DNA fragments distributed within the domain. The mixed PCR fragments were digested with 5 restriction enzymes (AluI, RsaI, DdeI, HaeIII, and BfuCI) and subjected to labeling by incorporation of Cy3-dCTP (GE HealthCare) and Cy5-dCTP (GE HealthCare) using random primer DNA labeling kit (Takara). A total of 100 ng FISH probes were suspended with hybridization buffer (50% formamide, 2× SCC, 5× Denhardt’s solution, and 10% dextran).

FISH experiments were performed as previously described (Kim et al., 2016a). Exponentially growing cells (approximately 1 x 10^8^ cells at OD_595_ = 0.5) were subjected to the IF procedure with anti-tubulin TAT1 antibody as described above. After washing the cells with PEMBAL buffer, they were washed 3 times with PEM buffer and fixed with 3% pFA at room temperature for 20 minutes. The fixed cells were incubated with 0.25 M glycine, washed twice with PEM buffer, treated with 0.1 N HCl at room temperature for 5 minutes, and washed again with PEM buffer. Cellular RNAs were digested with 50 µg RNase A (Takara Bio) in PEMBAL buffer at 37°C for 2 hours, and the cells were washed once with PEM buffer. FISH probes were denatured at 75°C for 15 minutes and mixed with the cells, and genomic DNA in the cells was denatured at 75°C for 5 minutes. Hybridization was performed at 40°C for 12 hours with a gentle rotation. The cells were washed 3 times at room temperature for 30 minutes with a gentle rotation. DNA was stained with 1 μg/mL DAPI in PBS buffer. The stained cells were mounted on a coverslip with the antifade solution. The fluorescent signals were detected using a DeltaVision deconvolution microscope. FISH experiments were repeated at least three times, and more than 30 cells were analyzed for respective samples unless otherwise noted.

### ChIP-seq

Fission yeast cells (approximately 5 x 10^8^ cells at OD_595_ = 0.5) were fixed with 3% pFA at 26°C for 30 minutes, washed twice with 1× PBS, and further fixed with 10 mM dimethyl adipimidate (Sigma Aldrich) at room temperature for 45 minutes. After two washes with 1× PBS, the cells were disrupted with glass beads using Mini-Beadbeater-16 (BioSpec Products) in ChIP lysis buffer [50 mM HEPES-KOH pH 7.5, 140 mM NaCl, 1 mM EDTA, 1% Triton X-100, 0.1% sodium deoxycholate, 0.1 mM PMSF, and EDTA-free Protease Inhibitor Cocktail (Roche)]. Chromatin DNA was sheared into 100–500 bp fragments by Bioruptor UCD-200 (Diagenode), and the soluble fraction was collected by centrifugation at 13,000 rpm for 15 minutes. A total of 50 μl ChIP lysis buffer containing 5 µl mouse monoclonal antibodies [anti-Pk (BioRad), anti-Myc (Takara Bio), and anti-Flag (Sigma)] and 15 µl Dynabeads protein G (Fisher Scientific) was added to the soluble fraction, and the bead suspension was incubated at 4°C for 2 hours with a gentle rotation. The beads were washed twice with ChIP lysis buffer, once with ChIP lysis buffer containing 0.65 M NaCl, once with washing buffer (10 mM Tris-HCl pH 8.0, 250 mM LiCl, 0.5% NP-40, 0.5% sodium deoxycholate, and 1 mM EDTA), and once with TE (10 mM Tris-HCl pH 8.0, and 1 mM EDTA) for 5 minutes each with a gentle rotation. The protein-DNA mixture was eluted by incubating the beads with TES (10 mM Tris-HCl pH 8.0, 1 mM EDTA, and 1% SDS) at 65°C for 30 minutes with harsh agitation at 1,000 rpm using ThermoMixer (Eppendorf) and subjected to reverse crosslinking at 68°C overnight in the presence of 0.5 M NaCl. DNA was purified using QIAquick PCR purification kit (Qiagen). The recovered DNA was subjected to library construction using NEBNext Ultra DNA library prep kit (NEB). The sequencing libraries were amplified by PCR using the NEBNext Ultra II Q5 Master Mix (NEB) with NEBNext Multiplex Oligos for Illumina (NEB). The libraries were sequenced by the NextSeq 500, NextSeq 2000, or NovaSeq 6000 (Illumina) to obtain paired-end reads.

### In situ Hi-C

Fission yeast cells (approximately 5 x 10^8^ cells at OD_595_ = 0.5) were fixed with 3% pFA at 26°C for 10 minutes and disrupted with glass beads using Mini-Beadbeater-16 (BioSpec Products) in lysis buffer [50 mM HEPES pH 7.5, 140 mM NaCl, 1 mM EDTA, 1% Triton X-100, 0.1% sodium deoxycholate, 0.1 mM PMSF, and EDTA-free Protease Inhibitor Cocktail (Roche)]. After washing with 1 ml PBS solution, the cells were further fixed with 3 mM disuccinimidyl glutarate (Fisher Scientific) in 2.5 ml PBS solution at 30°C for 40 minutes. The fixed cells were incubated at 62°C for 7 minutes with 0.1% SDS and subsequently with 1% Triton-X 100. The permeabilized cells were incubated with 25 units of MboI, HinfI, and MluCI at 37°C overnight. The pellet containing restriction enzyme-digested DNA fragments was treated with 150 μM biotin-14-dATP, dCTP, dGTP, and dTTP and incubated with 20 units of Klenow fragment (NEB) at 37°C for 45 minutes, followed by proximity ligation using 2,500 units of T4 DNA ligase (NEB) at room temperature for 4 hours with a gentle rotation. After ligation, the pellet was subjected to proteinase K treatment at 68°C overnight in the presence of 0.5 M NaCl. DNA was purified by phenol/chloroform extraction and ethanol precipitation. The purified DNA was sheared into 100– 500 bp fragments by Bioruptor UCD-200 (Diagenode). Biotin-labeled DNA was purified using Dynabeads MyOne Streptavidin T1 (Fisher Scientific). Sequencing adapters were ligated using the NEBNext Ultra II DNA Library Prep Kit for Illumina (NEB) while DNA was bound to the beads. Sequencing libraries were amplified by PCR using the NEBNext Ultra II Q5 Master Mix (NEB) with NEBNext Multiplex Oligos for Illumina (NEB). The libraries were sequenced by the NextSeq 500, NextSeq 2000, or NovaSeq 6000 (Illumina) to obtain paired-end reads.

### ChIP-seq data processing

ChIP-seq data were processed as described previously (Iwasaki et al., 2019; Tanaka et al., 2012) with modifications. ChIP-seq reads were aligned to the *S. pombe* genome (2018 version) using Bowtie2 (version 2.4.4). The ChIP-seq peaks were defined by MACS3 (version 3.0.1) with a subcommand “bdgpeakcall”. The option –c 5 or 4, –l 220, –g 300. –c was determined from the mean value of scores of each sample and the *p*-value < 0.1. For ChIP-seq data shown in **Figure 5**, enrichment scores were calculated by subtracting the read density of no tag control from that of the indicated sample.

### In situ Hi-C data processing

In situ Hi-C data were processed as described previously (Tanizawa et al., 2017). In situ Hi-C data were processed using the rfy_hic2 package (https://github.com/rafysta/rfy_hic2; Tanizawa et al., 2017). In brief, paired-end sequence reads were separately aligned to the *S. pombe* genome (2018 version) using an iterative alignment strategy with Bowtie2 (version 2.4.4). Redundant paired reads derived from PCR bias, reads aligned to repetitive sequences, and reads with low mapping quality (MapQ < 30) were removed. Read pairs with inward or outward orientations within 10 kb were excluded from the analysis, as they contain undigested products or self-ligation artifacts. Using the data of reads aligned in the same orientation, the number of correct Hi-C reads in inward and outward orientations was estimated. The fission yeast genome was divided into non-overlapping bins of 5 kb, 10 kb, and 20 kb to create raw contact matrices by counting the number of read pairs assigned to respective bin combinations. Hi-C biases in contact maps were corrected using the ICE method by repeating the normalization process 30 times (Imakaev et al., 2012).

## DATA AVAILABILITY

All relevant data that support the finding of this study are available from the authors upon request.

## SUPPLEMENTAL INFORMATION

Supplemental Information includes Supplemental Experimental Procedures, 6 figures, and Supplemental References.

## ACKNOWLEDGMENTS

We would like to thank the Yeast Genetic Resource Center (Osaka City University) for the pTN-TH7 cDNA library and Drs. Keith Gull and Jack Sunter for anti-tubulin TAT1 antibody, and the University of Oregon Genomics & Cell Characterization Core Facility for next-generation sequencing and microscopic analysis. We also thank Drs. Jeannie and Eric Selker for scientific and humane guidance, and Ms. Yuko Tsukamoto for technical assistance. This work was supported by the National Institutes of Health/National Institute of General Medical Sciences R01GM124195, National Institutes of Health/National Institute of Aging P01AG AG031862, JSPS KAKENHI grant JP20K23376, Takeda Science Foundation, Grant for Basic Science Research Projects from The Sumitomo Foundation, and The Mitsubishi Foundation (K.N.).

