## SUPPLEMENTAL INFORMATION for "A role for condensin-mediator interaction in mitotic chromosomal organization"

### SUPPLEMENTAL FIGURES

#### Figure S1

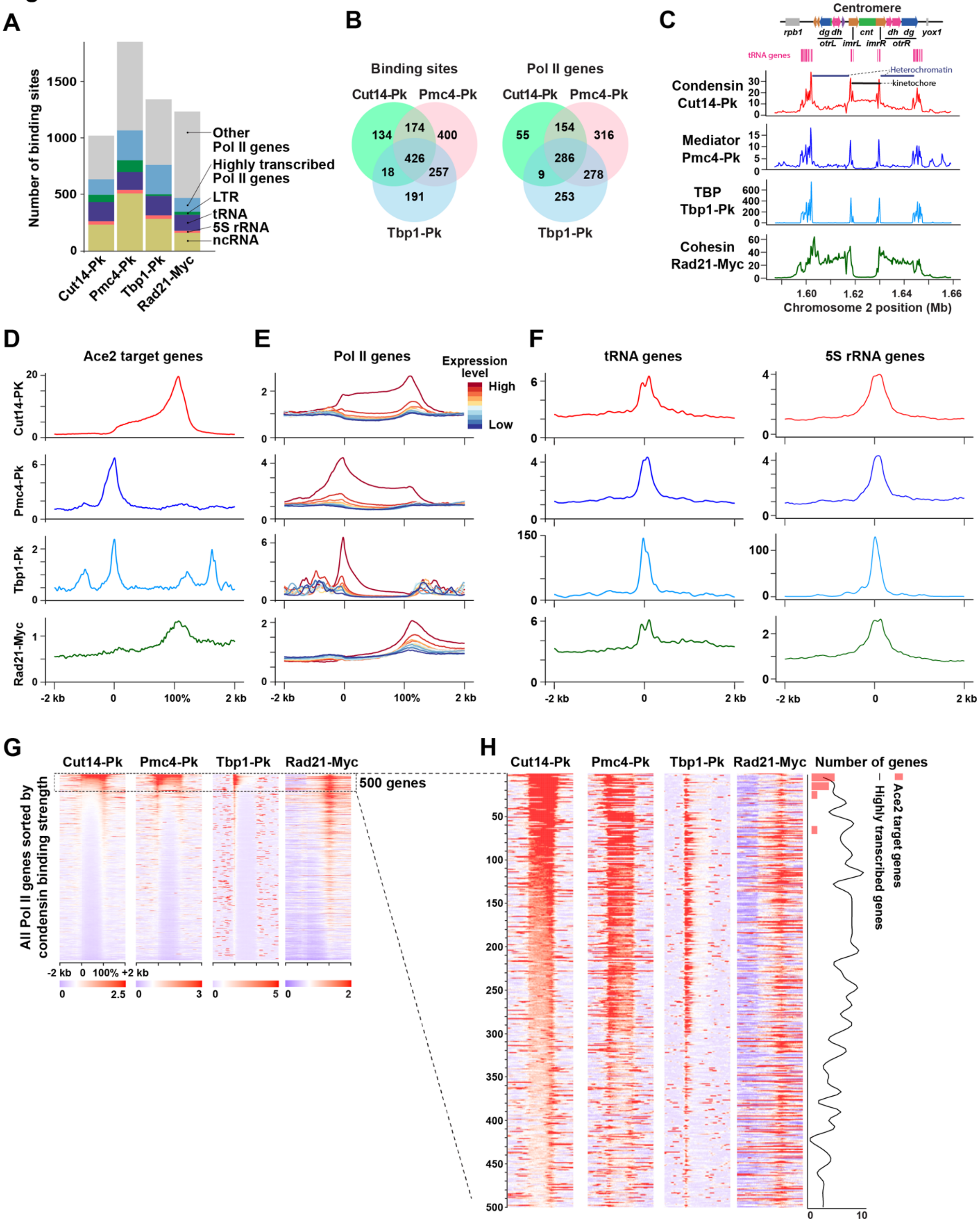

**Figure S1. Genome-wide distribution analysis of Cut14-Pk (condensin), Pmc4-Pk (mediator), Tbp1-Pk (TATA-box binding protein; TBP), and Rad21-Myc (cohesin)**

**(A)** Distributions of Cut14-Pk (condensin), Pmc4-Pk (mediator), Tbp1-Pk (TBP), and Rad21-Myc (cohesin) at the indicated genetic elements.

**(B)** Venn diagrams indicating the overlaps among Cut14-Pk, Pmc4-Pk, and Tbp1-Pk binding sites (left) and genes (right).

**(C)** Localization of Cut14-Pk, Pmc4-Pk, Tbp1-Pk, and Rad21-Myc at the centromere of chromosome 2.

**(D)** Average ChIP-seq enrichment of Cut14-Pk, Pmc4-Pk, Tbp1-Pk, and Rad21-Myc at Ace2 target genes.

**(E)** Correlation between transcription levels and binding of Cut14-Pk, Pmc4-Pk, Tbp1-Pk, and Rad21-Myc. Pol II genes (n=3405) were categorized into 10 groups based on their expression levels, and the average ChIP enrichment was plotted for the respective proteins.

**(F)** Average ChIP-seq enrichment of Cut14-Pk, Pmc4-Pk, Tbp1-Pk, and Rad21-Myc at tRNA genes (n=171; left) and 5S rRNA genes (n=33; right).

**(G)** Heatmaps showing ChIP-seq enrichment of Cut14-Pk, Pmc4-Pk, Tbp1-Pk, and Rad21-Myc at all Pol II genes. All Pol II genes were ranked by Cut14-Pk ChIP enrichment. Gene sizes from transcriptional initiation sites (0%) to termination sites (100%) were adjusted to the same length for data representation.

**(H)** Heatmaps showing ChIP-seq enrichment of Cut14-Pk, Pmc4-Pk, Tbp1-Pk, and Rad21-Myc at 500 Pol II genes with the highest Cut14-Pk enrichment. Distributions of Ace2 target genes and highly transcribed genes are shown in the right panel.

### Figure S2

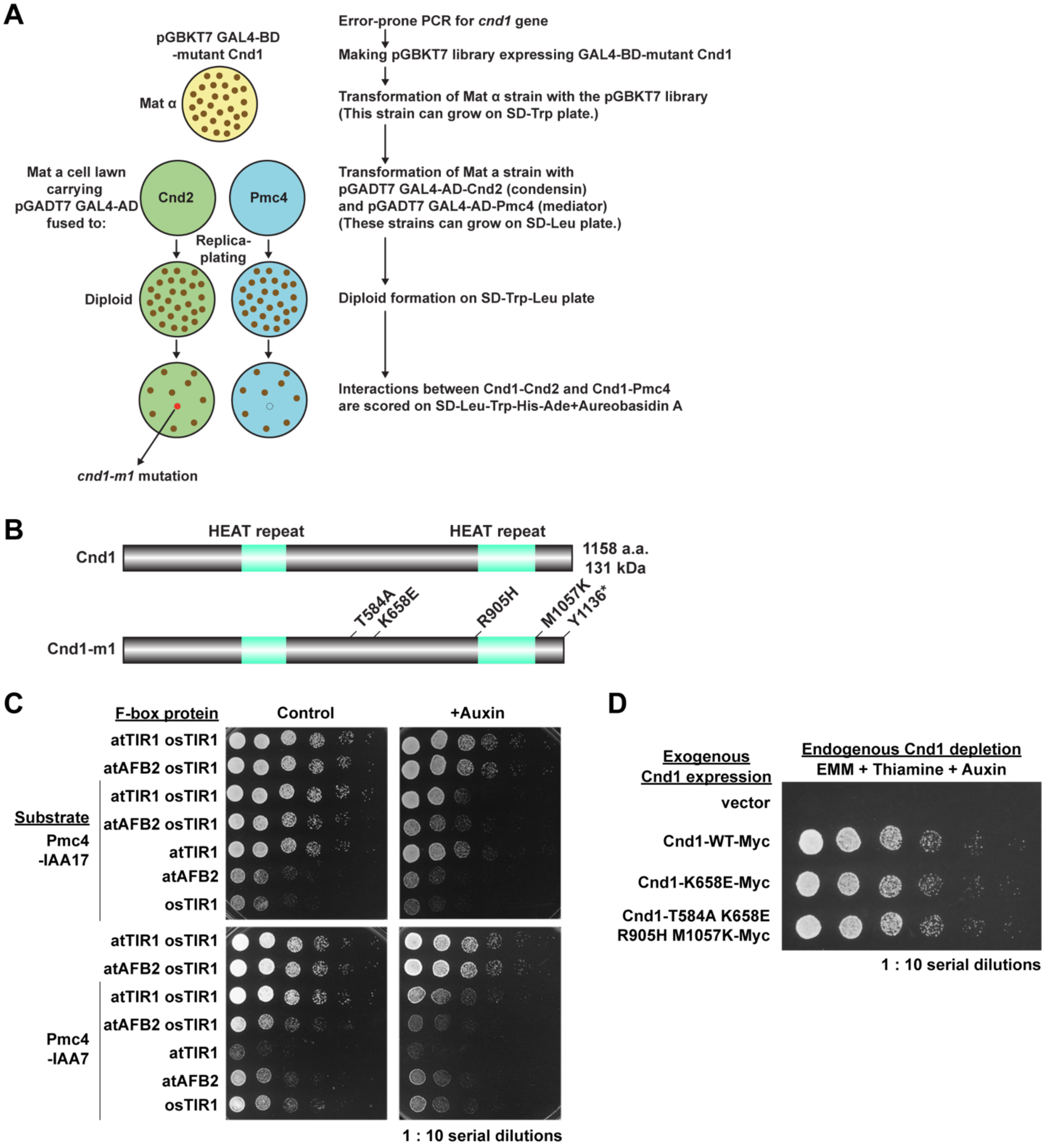


**Figure S2. Generation of *cnd1-m1* mutation and optimization of the AID system**

**(A)** Y2H-based random mutagenesis approach to identify the *cnd1-m1* mutation.

**(B)** The *cnd1-m1* mutation. Cnd1 consists of the two HEAT repeats, which are known to mediate interactions with kleisin subunits and are predicted to have functions related to DNA repair, kinetochore function, and ploidy maintenance (Piazza et al., 2014; Xu et al., 2015).

**(C)** Optimization of the AID system for fission yeast cells. The *ade6* locus expressing the F-box proteins (atTIR1, atAFB2, and/or osTIR1) was combined with the *pmc4* locus expressing Pmc4 fused to IAA7 or IAA17 degrons. Cells were cultured in YEA liquid medium without (control) and with auxin (Pmc4 depletion condition). Ten-fold serial dilutions of logarithmically growing cells were spotted on the YEA (control) and YEA + auxin plates.

**(D)** The suppression of Cnd1 depletion by exogenous expression of wild-type and mutant Cnd1 proteins. Cells were cultured in EMM liquid medium containing thiamine and auxin to induce transcriptional inhibition of the *cnd1* gene and post-translational degradation of endogenous Cnd1 proteins. The wild-type and mutant Cnd1 were expressed from plasmids. Ten-fold serial dilutions of logarithmically growing cells were spotted on the EMM + thiamine + auxin plate.

### Figure S3

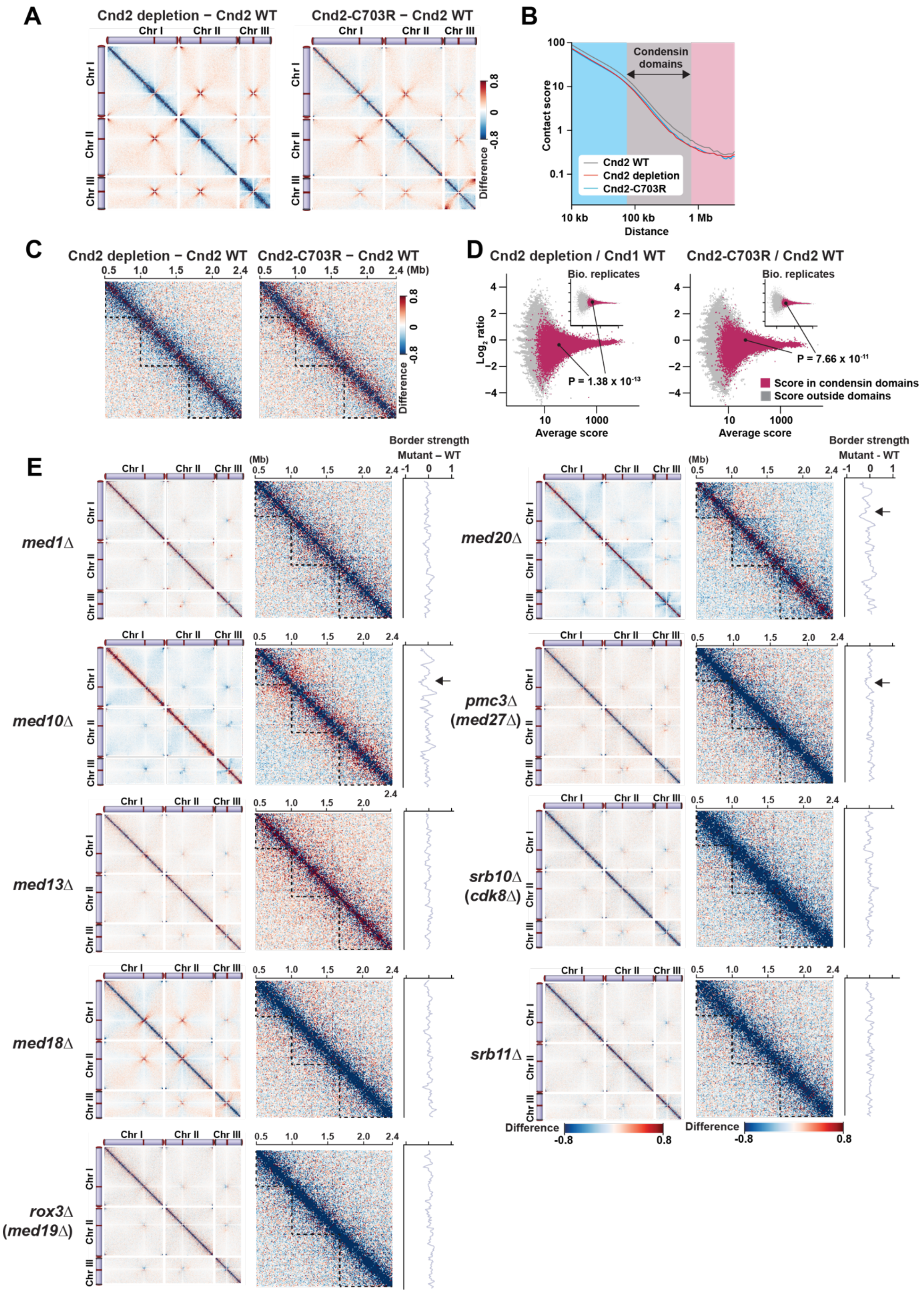

A
Cnd2 depletion − Cnd2 WT
Cnd2-C703R − Cnd2 WT
Chr I
Chr II
Chr III
Difference
B
Condensin domains
Contact score
Distance
Cnd2 WT
Cnd2 depletion
Cnd2-C703R
10 kb
100 kb
1 Mb
C
Cnd2 depletion − Cnd2 WT
Cnd2-C703R − Cnd2 WT
(Mb)
D
Cnd2 depletion / Cnd1 WT
Cnd2-C703R / Cnd2 WT
Bio. replicates
Log2 ratio
P = 1.38 x 10-13
P = 7.66 x 10-11
Average score
Score in condensin domains
Score outside domains
E
Border strength
Mutant – WT
med1Δ
med10Δ
med13Δ
med18Δ
rox3Δ (med19Δ)
med20Δ
pmc3Δ (med27Δ)
srb10Δ (cdk8Δ)
srb11Δ
Difference
-0.8
0.8

**Figure S3. Effects of *cnd2-C703R* mutation and mediator gene deletions on condensin domains**

**(A)** Difference maps showing the difference of contact scores between Cnd2 depletion and Cnd2 WT data (left) and between Cnd2-C703R and Cnd2 WT data (right). Cells were cultured in EMM liquid medium containing thiamine and auxin to induce transcriptional inhibition of the *cnd2* gene and post-translational degradation of endogenous Cnd2 proteins (Iwasaki et al., 2015). The Cnd2-C703R was expressed from the plasmid. Cnd2 depletion represents without exogenous Cnd2 expression. Asynchronous cells were used for this Hi-C analysis.

**(B)** Relations between contact scores and genomic distances.

**(C)** Enlarged difference maps. The genomic position is shown at the top. The dotted lines indicate previously defined condensin domains (Tanizawa et al., 2017).

**(D)** Statistical analysis on disruption of condensin domains by Cnd2 depletion and the Cnd2-C703R mutation. Averages and $\log_2$ ratios of contact scores between the indicated sample combinations were subjected to statistical analysis as described in **Figure 3E**, except that the two biological replicates from wild-type asynchronous cells were used as a control.

**(E)** Difference maps between the wild type and indicated mediator mutants for the entire genome (left) and the indicated 1.9 Mb region of chromosome I (right). The deletion mutants of the non-essential mediator genes (*med1*, *med10*, *med13*, *med18*, *rox3*, *med20*, *pmc3*, *srb10*, and *srb11*) and wild-type cells from asynchronous culture were subjected to in situ Hi-C analysis. The difference in the border strength scores (mediator mutant – WT) is shown on the right. Arrows indicate the position of the impaired chromatin boundary at the *eng1* locus in the mediator mutants.

### Figure S4

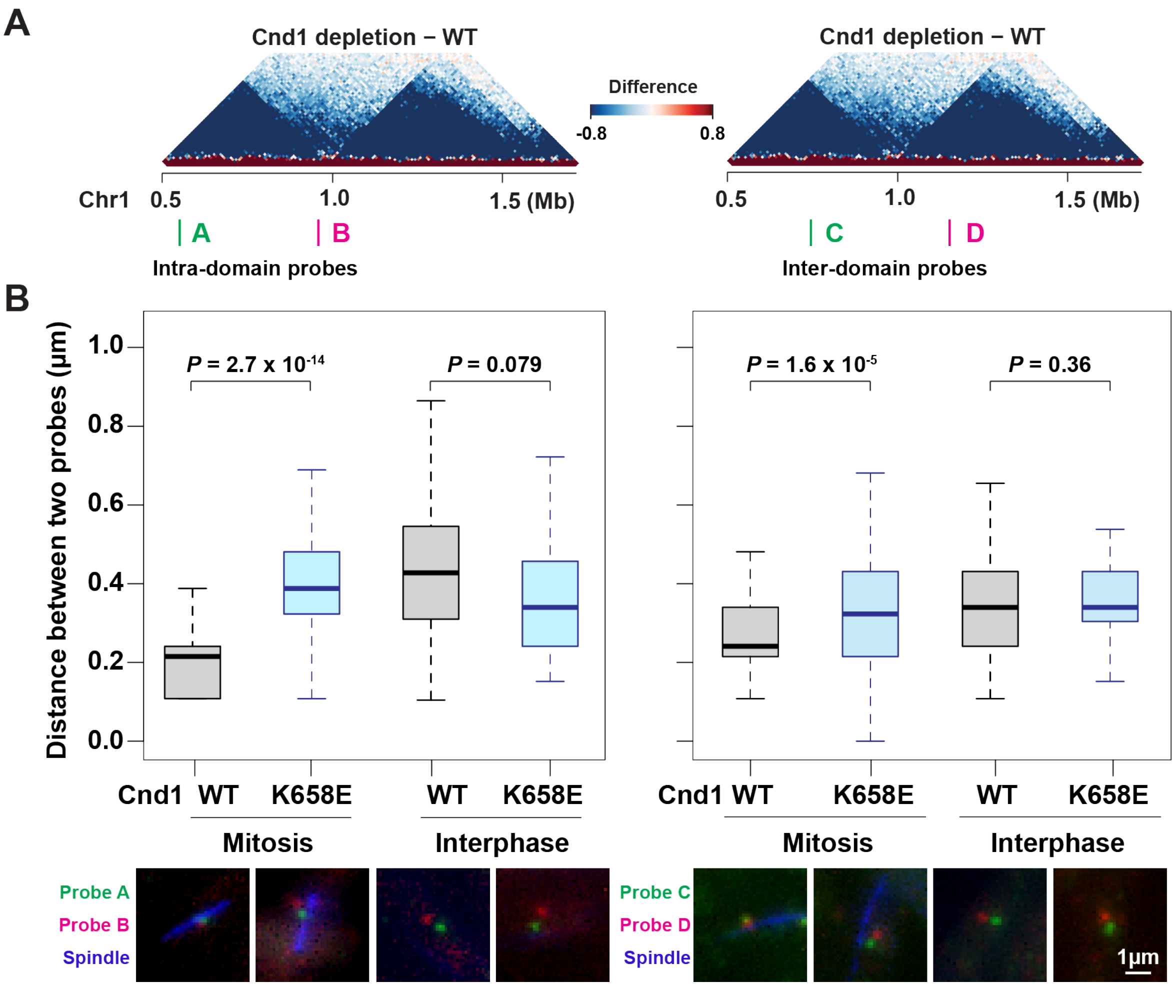


**Figure S4. FISH visualization of two pairs of genomic loci located within and spanning condensin domains**

**(A)** Positions of the FISH probes. Two pairs of probes were designed to visualize the two loci within the domain (left) and between the two domains (right).

**(B)** Quantification of distances between the two FISH probes within the domain (left) and between the domains (right). FISH data from interphase and mitotic cells with wild-type Cnd1 expression from the plasmid (Cnd1 WT), without exogenous Cnd1 expression (Cnd1 depletion), and with exogenous Cnd1-K658E expression (Cnd1-K658E) are summarized. Centers of FISH foci were defined as positions of the loci. The distance between the two loci was measured in more than 30 cells, and the same FISH experiments were triplicated. The central bar represents the median, with boxes indicating the upper and lower quartiles. Whiskers extend to the data points of no more than 1.5× the interquartile range from the box. Typical FISH-IF images visualizing the genomic loci (red and green) and spindle (blue) are shown at the bottom.

### Figure S5

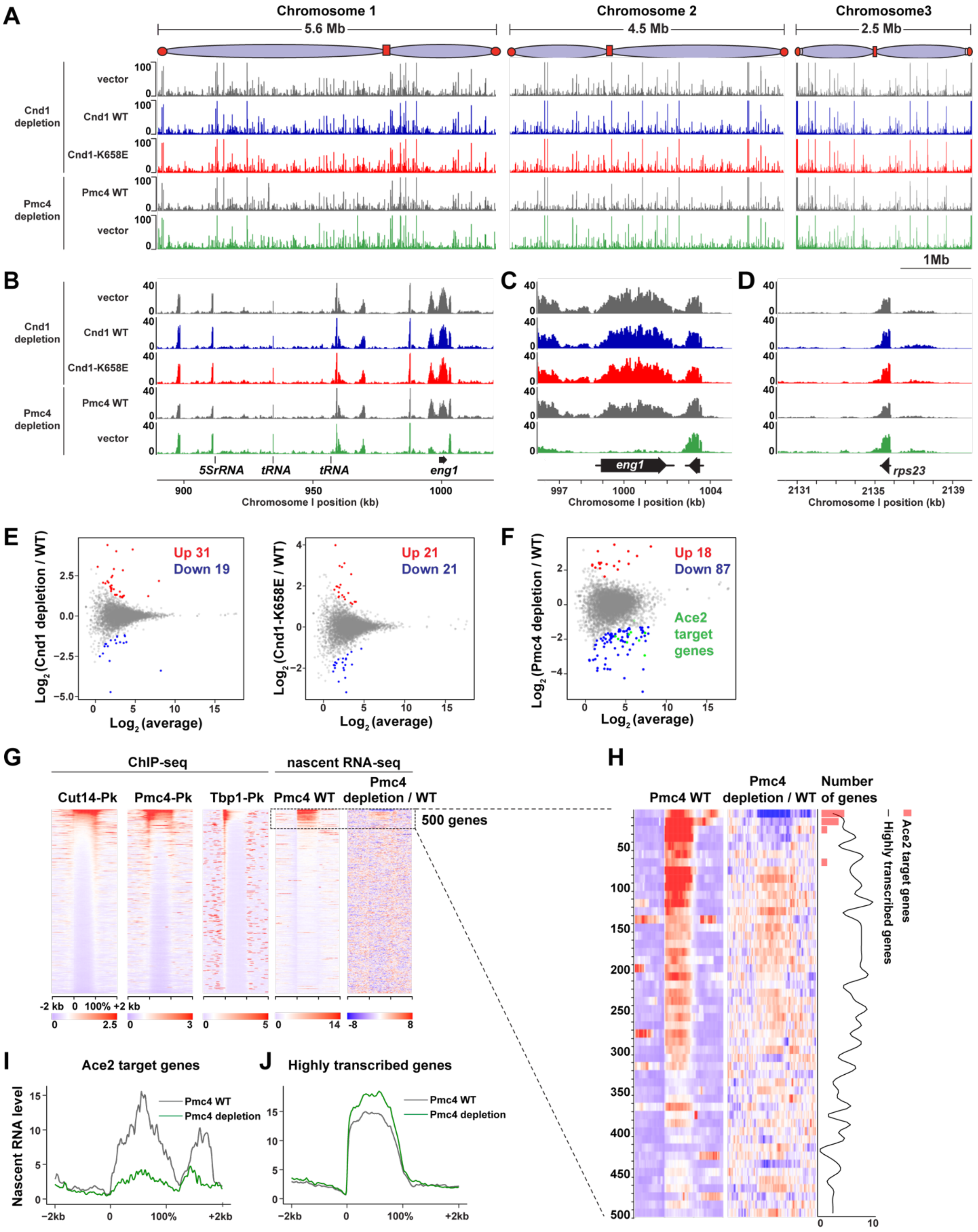


**Figure S5. Effects of Pmc4 depletion on gene expression**

**(A)** Nascent RNA profiles in the indicated conditions were determined by nascent RNA-seq. The endogenous Cnd1 expression was depleted, and the exogenous Cnd1 proteins (WT and K658E) were provided from plasmids. The vector represents Cnd1 depletion without the exogenous Cnd1. For Pmc4 depletion, the endogenous Pmc4 was degraded by the AID system, and the exogenous Pmc4 protein was provided from the plasmid (Pmc4 WT). The vector sample represents Pmc4 depletion without the exogenous Pmc4 expression.

**(B-D)** Nascent RNAs at the 130 kb genomic region of chromosome I **(B)**, the *eng1* Ace2 target gene locus **(C),** and the *rps23* highly transcribed gene locus **(D)**.

**(E)** Scatter plots showing nascent RNA ratios (Y-axis) and average nascent RNA levels (X-axis) of respective genes between Cnd1 depletion and Cnd1 WT samples (left) and between Cnd1-K658E and Cnd1 WT samples (right).

**(F)** Scatter plot showing nascent RNA ratios samples (Y-axis) and average nascent RNA levels (X-axis) of respective genes between Pmc4 depletion and Pmc4 WT samples. Eight Ace2 target genes (total 9) were significantly down-regulated (green).

**(G)** Heatmaps showing ChIP-seq enrichment of Cut14-Pk, Pmc4-Pk, Tbp1-Pk, nascent RNA levels, and nascent RNA ratios between Pmc4 depletion and Pmc4 WT samples. All Pol II genes were ranked by Cut14-Pk ChIP enrichment.

**(H)** Enlarged heatmaps for 500 Pol II genes with the highest Cut14-Pk enrichment. Distributions of Ace2 target genes and highly transcribed genes are shown in the right panel.

(**I** and **J**) Average nascent RNA levels in Pmc4 depletion and Pmc4 WT samples at Ace2 target genes (**I**) and highly transcribed genes **(J)**.

### Figure S6

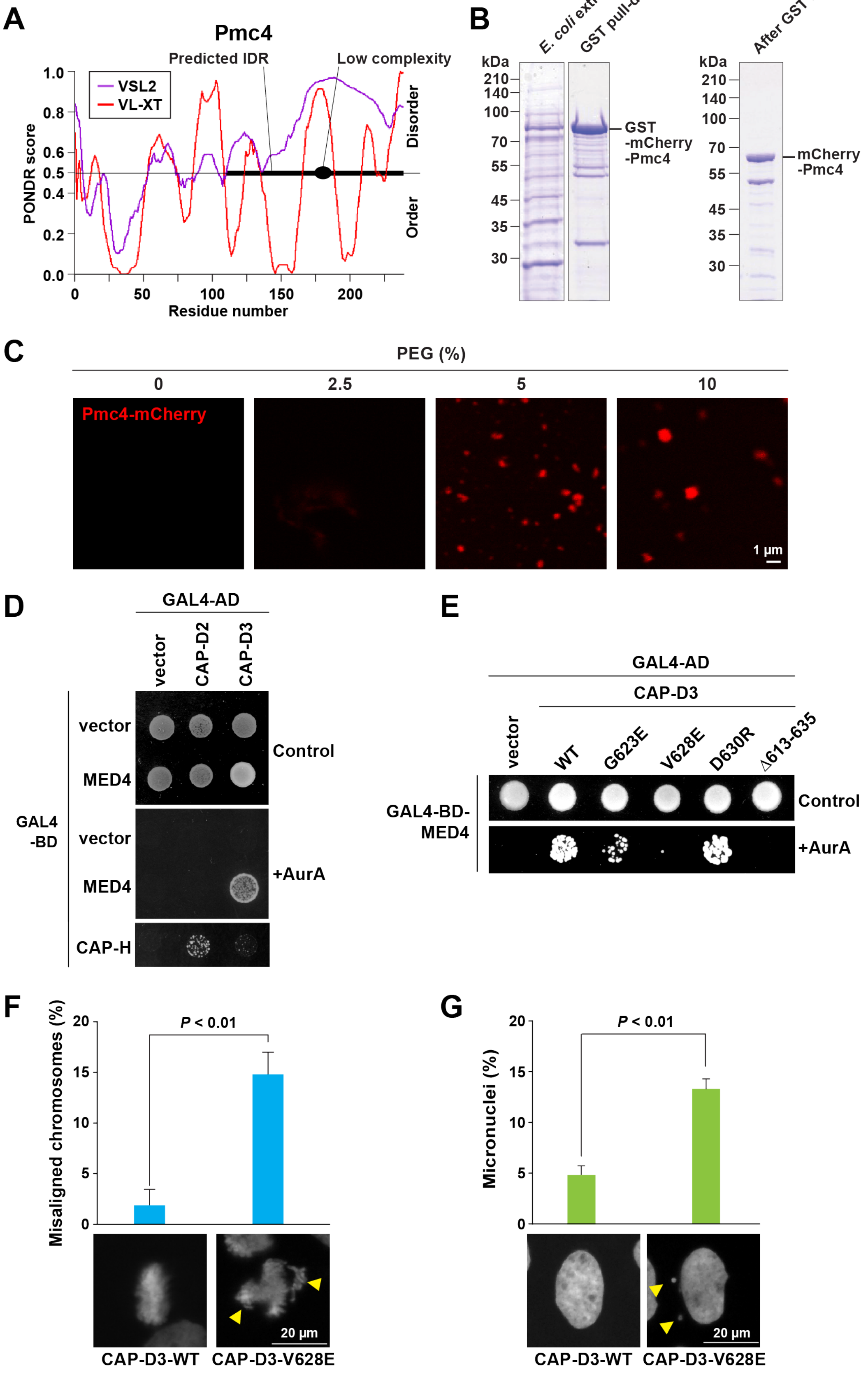

A
Pmc4
Predicted IDR
Low complexity
VSL2
VL-XT
PONDR score
Disorder
Order
Residue number
B
E. coli extract
GST pull-down
After GST removal
kDa
GST
-mCherry
-Pmc4
mCherry
-Pmc4
C
PEG (%)
Pmc4-mCherry
1 µm
D
GAL4-AD
vector
CAP-D2
CAP-D3
GAL4
-BD
vector
MED4
CAP-H
Control
+AurA
E
GAL4-AD
CAP-D3
vector
WT
G623E
V628E
D630R
Δ613-635
GAL4-BD-
MED4
Control
+AurA
F
Misaligned chromosomes (%)
P < 0.01
20 µm
CAP-D3-WT
CAP-D3-V628E
G
Micronuclei (%)
P < 0.01
20 µm
CAP-D3-WT
CAP-D3-V628E

**Figure S6. Potential involvement of Pmc4 in phase separation**

**(A)** PONDR prediction of IDRs. VSL2 and VL-XP prediction models were employed. The predicted IDR is shown as a thick line (Romero et al., 2001).

**(B)** Purification of fission yeast Pmc4 proteins from *E. coli* cells. GST-Pmc4-mCherry proteins were expressed in *E. coli* cells and purified by GST-pull down. GST was removed by PreScission Protease treatment.

**(C)** Pmc4-mCherry tested for droplet formation in varying concentrations of PEG.

**(D)** Y2H interaction detected between CAP-D3 and MED4. MED4 (human Pmc4) fused to GAL4-BD and CAP-D2 (human Cnd1 in condensin I) and CAP-D3 (human Cnd1 in condensin II) fused to GAL4-AD were expressed in budding yeast cells. The Y2H interaction between CAP-H (human Cnd2 in condensin I) and CAP-D2 is shown as a positive control, indicating that CAP-D2 is expressed in budding yeast cells and interacts with CAP-H as expected.

**(E)** Y2H assay testing the MED4 interaction with mutant CAP-D3 proteins. Based on sequence alignment between Cnd1 and CAP-D3, point mutations were introduced near amino acid residues within CAP-D3 aligned with the lysine 658 residue of Cnd1. CAP-D3-V628E mutation disrupted the Y2H interaction between CAP-D3 and Med4.

**(F)** Frequencies of misaligned chromosomes in human U2OS cells expressing wild-type or mutant CAP-D3 proteins. Arrowheads indicate misaligned chromosomes during metaphase in U2OS cells expressing CAP-D3-V628E.

**(G)** Frequencies of micronuclei detected in human U2OS cells expressing wild-type or mutant CAP-D3 proteins. Arrowheads indicate micronuclei observed in U2OS cells expressing CAP-D3-V628E.

In panels **(F)** and **(G)**, more than 30 mitotic cells were examined, and results from three biological replicates are shown. Data are represented as mean ± SD.

### EXTENDED EXPERIMENTAL PROCEDURES

#### Nascent RNA labeling followed by deep sequencing (nascent RNA-seq)

The nascent RNA detection protocol used in this study was based on the previously reported precision run-on sequencing (PRO-seq) approach (Mahat et al., 2016) with some modifications. A total of $1\times10^8$ cells were permeabilized in 0.5% N-Lauroylsarcosine sodium salt (sarkosyl; Sigma) on ice for 20 minutes, and a nuclear run-on reaction was carried out at 30°C for 5 minutes in a reaction buffer [20 mM Tris-HCl pH 7.7, 200 mM KCl, 5 mM $MgCl_2$, 1.25 mM dithiothreitol, 0.4 U/µL SUPERase-In RNase Inhibitor (Fisher), 0.5% sarkosyl, 12.5 µM biotin-11-CTP (Enzo Life Sciences), 12.5 µM biotin-11-UTP (Biotium), 62.5 µM ATP (Fisher), and 62.5 µM GTP (Fisher)] to label newly transcribed RNA with biotin. The cells after the pulse-labeling reaction were disrupted with Mini-Beadbeater-16 in the presence of AES buffer [50 mM sodium acetate pH 5.5 (Fisher), 10 mM EDTA, and 1% SDS], neutral Phenol Chloroform Isoamyl alcohol (PCI), and glass beads. Nucleic acids were purified by additional neutral PCI extraction followed by ethanol precipitation. RNA was heat-denatured at 65°C for 1 minute and fragmented by 0.2 N sodium hydroxide (NaOH; Fisher) treatment on ice for 10 minutes. After neutralization with 0.5 M Tris-HCl pH 6.8, the fragmented RNA was purified with RNeasy Mini Kit (Qiagen), and residual DNA was removed by 0.05 U/µL RQ1 RNase-Free DNase (Promega) treatment at 37°C for 30 minutes followed by neutral PCI extraction and ethanol precipitation. The biotinylated RNA was selectively pulled down using Dynabeads MyOne Streptavidin T1 magnetic beads (Fisher) at 25°C for 15 minutes, and first- and second-strand cDNA syntheses, end repair, and adaptor ligation reactions were successively performed using NEBNext Ultra II RNA Library Prep Kit for Illumina (NEB), according to the manufacturer instruction. The adaptor-ligated cDNA was amplified using NEBNext Ultra II Q5 Master Mix and NEBNext Multiplex Oligos for Illumina.

The PCR-amplified libraries were purified with 0.9× volume of AMPure XP before sequencing runs with NextSeq 500 or NextSeq 2000 to obtain paired-end reads.

**Nascent RNA-seq analysis**

Sequenced reads were aligned to the *Schizosaccharomyces pombe* genome (2018 version) using STAR (version 2.7.6) (Dobin et al., 2013). Reads assigned to exons were estimated by the RSEM program (version 1.3.3) (Li and Dewey, 2011). Normalization of read numbers and definition of differentially expressed genes between WT and mutants were calculated using edgeR program version 4.0.16 (Robinson et al., 2010). The dispersion values for Cnd1 expression (vector, WT, and K658E mutant) and Pmc4 (WT and depletion) were 0.017 and 0.037, respectively. FDR < 0.05 was used as a threshold to define significantly up- or down-regulated genes.

**Pmc4 purification from *E. coli***

To induce GST-mCherry-Pmc4 protein expressions, BL21 *Escherichia coli* cells carrying the pGEX6P-mCherry-Pmc4 plasmid were cultured in LB medium supplemented with 10 μM IPTG (Wako) at 16 °C for 20 hours. Cellular proteins were extracted with extraction buffer [50 mM Tris-HCl (pH 8.0), 500 mM NaCl, 10 mM β-mercaptoethanol, and 10% glycerol], and affinity purification was performed using GST-accept resin (Nakarai). The bound protein was eluted with glutathione buffer [10 mM reduced glutathione (Wako), 50 mM Tris-HCl, 500 mM NaCl, 10 mM β-mercaptoethanol, and 10% glycerol]. Purified proteins were treated with precision protease at 4 °C for 16 hours, followed by the desalting step using a HiTrap Q HP column (GE Healthcare) and the GST removal step using GST-accept. The buffer was exchanged with LLPS buffer (50 mM Tris-HCl, 100 mM NaCl, and 10 mM β-mercaptoethanol) using VivaSpin 6 Centrifugal

Concentrators (Sartorius). The purified protein solution was mixed with PEG3350 buffer (100μM Bis-Tris pH6.0) containing the indicated concentrations of PEG3350 at a 1:1 ratio for droplet observation under the Leica STELLARIS 5 confocal microscope equipped with an HC PL APO CS2 63x/1.40 OIL objective lens (Leica). Images were captured and analyzed using LAS X 4.4.0.24861 software (Leica).

**Human cell lines and culture conditions**

Plasmids expressing eGFP-CAP-D3-WT or eGFP-CAP-D3-V628E under the control of the TRE3GS doxycycline-inducible promoter were transfected into U2OS cells together with pTORA14_AAVS using Lipofectamine LTX Reagent (Fisher Scientific). To establish stable cell lines, the cells were cultured with DMEM containing 10% FBS and 10 μg/mL puromycin for 14 days, with refreshing the medium every 3 days.

**DAPI staining with human cells**

To observe the mitotic chromosomes and micronuclei, U2OS cells stably maintaining eGFP constructs (eGFP-CAP-D3-WT and eGFP-CAP-D3-V628E) were cultured on cover glasses, and 2 μg/mL doxycycline was added 24 hours before fixation. The cells were fixed with 4% pFA in PBS at 37ºC for 10 minutes. After treatment with 0.15% Triton-X in PBS for 2 minutes at room temperature, the cells were washed with PBS and mounted on slide glasses using an Antifade mounting medium (Vectashield) containing 1 μg/mL DAPI. The images were acquired using a confocal microscope (Olympus, FV1000) with a 60× lens.
